# A role for the auxin precursor anthranilic acid in root gravitropism via regulation of PIN-FORMED protein polarity and relocalization in *Arabidopsis*

**DOI:** 10.1101/422733

**Authors:** Siamsa M. Doyle, Adeline Rigal, Peter Grones, Michal Karady, Deepak K. Barange, Mateusz Majda, Barbora Pařízková, Michael Karampelias, Marta Zwiewka, Aleš Pěnčik, Fredrik Almqvist, Karin Ljung, Ondřej Novák, Stéphanie Robert

## Abstract

- Distribution of auxin within plant tissues is of great importance for developmental plasticity, including root gravitropic growth. Auxin flow is directed by the subcellular polar distribution and dynamic relocalization of auxin transporters such as the PIN-FORMED (PIN) efflux carriers, which can be influenced by the main natural plant auxin indole-3-acetic acid (IAA). Anthranilic acid (AA) is an important early precursor of IAA and previously published studies with AA analogues suggested that AA may also regulate PIN localization.
- Using *Arabidopsis thaliana* as a model species, we studied an AA-deficient mutant displaying agravitropic root growth, treated seedlings with AA and AA analogues and transformed lines to over-produce AA while inhibiting its conversion to downstream IAA precursors.
- We showed that AA rescues root gravitropic growth in the AA-deficient mutant at concentrations that do not rescue IAA levels. Overproduction of AA affects root gravitropism without affecting IAA levels. Treatments with, or deficiency in, AA result in defects in PIN polarity and gravistimulus-induced PIN relocalization in root cells.
- Our results reveal a previously unknown role for AA in the regulation of PIN subcellular localization and dynamics involved in root gravitropism, which is independent of its better-known role in IAA biosynthesis.

## Introduction

Auxin distribution in controlled concentration gradients within certain tissues plays an important role in regulating the dynamically plastic growth and development of plants (Vanneste & Friml, 2009). An intense research effort has revealed many of the complex mechanisms by which plasma membrane-localized auxin carrier proteins are polarly distributed in order to direct the flow of auxin in plant tissues and maintain these gradients (reviewed by Luschnig & Vert, 2014; and Naramoto, 2017). These proteins, including the well-studied PIN-FORMED (PIN) auxin efflux carriers, are remarkably dynamic in that they rapidly relocalize within the cell in response to signals, resulting in changes in their polarity. This dynamic responsiveness, which is facilitated by vesicular cycling and complex endomembrane trafficking pathways, is essential for altering the direction and strength of cell-to-cell auxin flow and redistributing auxin in response to external cues, thereby regulating cell and tissue growth and plasticity.

Root development in *Arabidopsis thaliana* has received particular attention as a model system demonstrating the importance of auxin gradients for plant development (Clark *et al*., 2014). Mutations affecting auxin transporters often disturb root gravitropism and specific PIN proteins within the root tip have been shown to relocalize in response to changes in the gravity vector, leading to altered auxin flow and consequently, organ growth adjustment (reviewed by Geisler *et al*., 2014). In the root columella, the cellular relocalization of PIN3 and PIN7 plays an important role in root gravitropic growth responses. While these proteins are generally apolar in columella cells, they redistribute toward the downward-facing plasma membranes upon horizontal reorientation of the root (Friml *et al*., 2002b; Kleine-Vehn *et al*., 2010), which is presumed to redirect the flow of auxin within the columella, thus contributing to auxin accumulation at the lower root side. PIN2 is also involved the root gravitropic response. Displaying shootward (apical) polarity within root epidermal cells (Müller *et al*., 1998), PIN2 transports auxin upward, balancing the correct auxin maximum required in the root apical meristem for root development (Adamowski & Friml, 2015). However, in the case of a horizontal reorientation of the root, PIN2 is rapidly redistributed from the plasma membranes to the vacuoles within epidermal cells at the upper organ side (Abas *et al*., 2006; Kleine-Vehn *et al*., 2008). This results in accumulation of auxin and consequent inhibition of cell elongation at the lower root side, contributing toward the root tip bending downward.

In our previous work, we employed a chemical biology approach, whereby we isolated and characterized small synthetic molecules selectively altering the polarity of specific PIN proteins, to dissect the trafficking pathways involved in regulating their localization (Doyle *et al*., 2015a). This approach led us to identify a potential role for the endogenous compound anthranilic acid (AA) in PIN polarity regulation, which we investigated in the current study. AA is an important early precursor of the main natural plant auxin indole-3-acetic acid (IAA) (Maeda & Dudareva, 2012) and as auxin itself has been shown to regulate PIN polarity in a feedback mechanism to control its own flow (Paciorek *et al*., 2005), we hypothesized that AA may play a similar regulatory role. Herein, using *Arabidopsis* root gravitropism as a model system for auxin-regulated plastic growth, we provide strong evidence in favor of this hypothesis. Ultimately, we reveal a previously unknown role for AA in the regulation of PIN polarity and relocalization required for root gravitropic responses and furthermore, we show that this role of AA is distinct from its well-known role in IAA biosynthesis.

## Materials and Methods

### Plant material and growth conditions

*Arabidopsis thaliana* was grown vertically on ½ strength Murashige and Skoog (MS) medium at pH 5.6 with 1% sucrose, 0.05% 2-(*N*-morpholino)ethanesulfonic acid (MES) and 0.7% plant agar for 5 or 9 d at 22 °C, on a 16 h : 8 h light : dark photoperiod. The Columbia-0 (Col-0) accession was used as wild type (WT). See Table S1 for the previously published *Arabidopsis* lines used and Table S2 for the genotyping primers used. All mutants/marker lines in the *wei2wei7* background were generated in this study by crossing (Methods S1). For generation of *35S::ASA1* (*35S::WEI2*) and *XVE::amiRNA-PAT1* lines and root growth measurements, see Methods S1.

### Chemical treatments and IAA metabolite analysis

Stock solutions of Endosidin 8 (ES8) (ID 6444878; ChemBridge, San Diego, CA, USA), AA (Sigma-Aldrich, St. Louis, MO, USA), ES8.7 (ID 6437223; ChemBridge) and ES8.7-Trp (Methods S2) were made in dimethyl sulfoxide (DMSO) and diluted in liquid medium for short-term (2 h) or growth medium for long-term (5 or 9 d) treatments, in which case seeds were directly sown on chemical-supplemented medium. Equal volumes of solvent were used as mock treatments for controls. For quantification of endogenous IAA and its metabolites, 20-30 whole seedlings per sample were flash-frozen in liquid nitrogen and c. 20 mg of ground tissue was collected per sample. Extraction and analysis were performed according to Novák *et al*. (2012) (Methods S2). See Methods S3 for compound degradation analysis.

### qPCR, immunolocalization and confocal microscopy

Quantitative real-time PCR (qPCR) was performed as described previously (Doyle *et al*., 2015a) (Methods S4). See Table S2 for the qPCR primers used. For β-glucuronidase (GUS) staining, see Methods S4. Immunolocalization was performed as described previously, using an InsituPro Vsi (Intavis Bioanalytical Instruments AG, Köln, Germany) (Doyle *et al*., 2015a). Primary antibodies used were anti-PIN1 at 1:500 (Nottingham Arabidopsis Stock Centre; NASC), anti-PIN3 at 1:150 (NASC), anti-PIN4 at 1:400 (NASC) and anti-PIN7 at 1:600 (Methods S4). Secondary antibodies used were Cy3-conjugated anti-rabbit and anti-sheep at 1:400 and 1:250, respectively (Jackson ImmunoResearch, Cambridgeshire, UK). Confocal laser scanning microscopy was performed using a Zeiss (Oberkochen, Germany) LSM 780 confocal microscope (see Methods S5 for microscopy image quantifications).

### Statistical analyses

For all experiments, at least three biological replicates were performed and always on different days. Occassionally, extra biological replicates were performed, due to poor growth of *wei2wei7* that sometimes resulted in a low number of seedlings or quantifiable roots in certain replicates. Unless indicated otherwise, Wilcoxon rank sum (Mann–Whitney U) tests or Student’s t-tests were performed on full, raw datasets of nonparametric or parametric data, respectively, to determine statistically significant differences and the means of the biological replicates are displayed on charts.

## Results

### AA rescues root gravitropic growth and length differently in an AA-deficient mutant

Using a chemical biology approach, we previously isolated the small synthetic molecule Endosidin 8 (ES8), which disturbs the polarity of selective PIN proteins in *Arabidopsis* roots, leading to altered auxin distribution patterns and defective root growth (Doyle *et al*., 2015a). Intriguingly, the chemical structure of ES8 reveals that this molecule is an analog of the endogenous plant compound AA (Fig. 1a), a precursor of tryptophan (Trp), the main precursor of the predominant plant auxin IAA (Ljung, 2013; Zhao, 2014). This prompted us to question whether endogenous AA might play a role in growth and development of the root. We therefore investigated a loss-of-function *Arabidopsis* mutant in both *ANTHRANILATE SYNTHASE SUBUNIT ALPHA1* (*ASA1*, also known as *WEAK ETHYLENE INSENSITIVE2*, *WEI2*) and *ANTHRANILATE SYNTHASE SUBUNIT BETA1* (*ASB1*, also known as *WEI7*). In the double mutant, *wei2wei7* (Stepanova *et al*., 2005; Ikeda *et al*., 2009), the AA level is presumed to be reduced. To confirm this, we analyzed the levels of several IAA precursors/catabolites, revealing that AA content was indeed significantly reduced in *wei2wei7* compared to the wild type (WT) Columbia-0 (Col-0), as were the levels of the IAA precursors Trp, indole-3-acetonitrile (IAN) and indole-3-acetimide (IAM) and IAA itself (Fig. S1), most likely due to the decreased AA content. However, neither the IAA precursor tryptamine (Tra) nor catabolite 2-oxoindole-3-acetic acid (oxIAA) showed altered content in the mutant compared to the WT (Fig. S1).

**Fig. 1.**
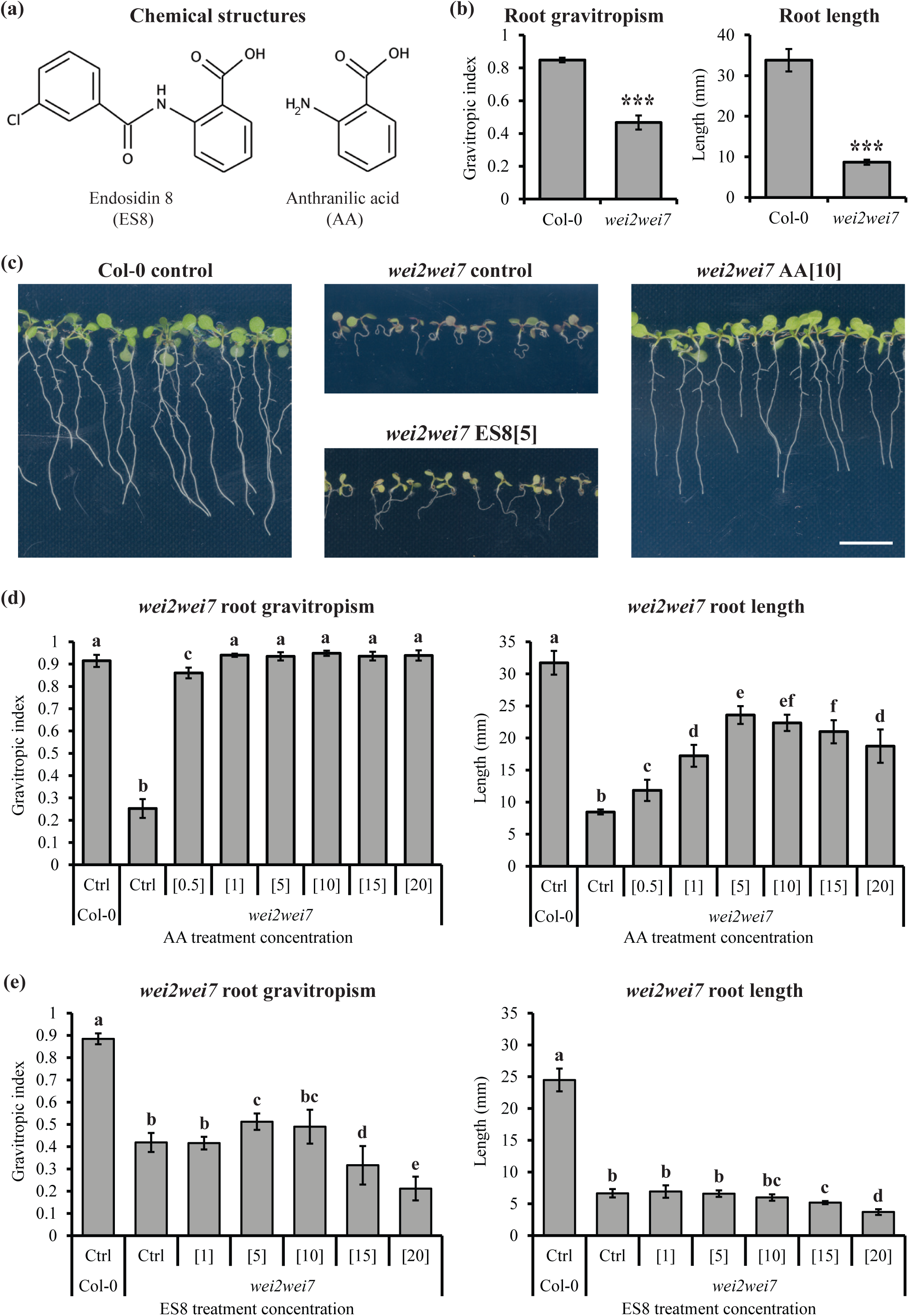
Anthranilic acid (AA) rescues root gravitropic growth and length differently in an AA-deficient mutant. (a) Chemical structures of Endosidin 8 (ES8) and AA. (b) Root gravitropic index and length in 9-d-old *Arabidopsis thaliana* Columbia-0 (Col-0) and *wei2wei7* seedlings. (c) Representative images of 9-d-old Col-0 and *wei2wei7* seedlings grown on treatment-supplemented medium. Scale bar represents 1 cm. (d-e) Root gravitropic index and length in 9-d-old seedlings of Col-0 and *wei2wei7* grown on mock-treated control medium (Ctrl) and *wei2wei7* grown on medium supplemented with a range of concentrations of AA (d) or ES8 (e). Asterisks indicate samples significantly different from Col-0 (***, *P*<0.001) (b) and different letters indicate significant differences (*P*<0.05) (d and e). Error bars indicate standard error of the mean of the biological replicates. Values in square brackets indicate treatment concentrations in µM. *n* = 25 seedlings per sample per each of four (b), three (d) or six (e) biological replicates.

We were interested in the strong agravitropic and short phenotypes of *wei2wei7* roots compared to Col-0 seedlings (Fig. 1b,c), considering that ES8 treatment reduces both gravitropic root growth and root length in Col-0 (Doyle *et al*., 2015a). To investigate AA-mediated rescue of these root phenotypes in *wei2wei7*, we performed long-term AA treatments by growing WT and mutant seedlings on medium supplemented with a range of AA concentrations. In Col-0, none of the tested concentrations affected root gravitropic growth, while concentrations of 10 µM or more decreased root length in a dose-dependent manner (Fig. S2a), possibly due to increased IAA biosynthesis. As expected, in *wei2wei7* both root gravitropic growth and length were rescued by AA (Fig. 1c,d), however we observed a striking difference between the AA rescue patterns of these two root phenotypes. While root gravitropic growth in the mutant was almost fully rescued to the WT level at all AA concentrations applied, root length rescue was only partial and was concentration-dependent, with maximal rescue at 5 µM (Fig. 1d). We hypothesized that these different rescue patterns in *wei2wei7* might reflect two different roles of AA, one known role in auxin biosynthesis and a distinct, as yet unknown role in regulating auxin distribution, considering that ES8 disturbs auxin distribution patterns in the root (Doyle *et al*., 2015a).

We next investigated whether ES8, as an analog of AA, could rescue either root gravitropic growth or length in *wei2wei7*. While long-term treatments with ES8 decreased both root gravitropic growth and length in a dose-dependent manner in Col-0 (Fig. S2b), only the highest ES8 concentrations (15 and 20 µM) decreased root gravitropic and length in *wei2wei7* (Fig. 1e). Moreover, while root length was not rescued in the mutant at any ES8 concentration, 5 µM ES8 partially rescued the root gravitropic phenotype of the mutant (Fig. 1c,e). The partial root gravitropic rescue of *wei2wei7* by ES8 without any effect on root length suggest, considering that ES8 is known to affect auxin distribution in the root (Doyle *et al*., 2015a), that the root gravitropic rescue of *wei2wei7* by AA may occur via a previously unknown role of AA in auxin distibution.

To further test our hypothesis, we used another analog of both ES8 and AA - ES8 analog no. 7 (ES8.7; Fig. S3a) and its analog ES8.7-Trp, in which the AA was exchanged for a Trp (Fig. S3b). In Col-0, long-term ES8.7 treatment revealed a similar but weaker effect than ES8 on dose-dependent reduction of root gravitropic growth and length (Fig. S3c). ES8.7 rescued root gravitropic growth in *wei2wei7* at a range of concentrations from 1 to 15 µM, with almost no effects on root length (Fig. S3a,d). Moreover, ES8.7-Trp did not rescue root gravitropic growth or length at any concentration in either Col-0 (Fig. S3e) or *wei2wei7* (Fig. S3b,f), strongly suggesting that it is the AA part of ES8 and ES8.7 that rescues gravitropic growth of *wei2wei7* roots. Together, these results suggest that a potential role for AA in auxin distribution may regulate root gravitropic growth, while the well-known role of AA in auxin biosynthesis may be more important for root length regulation.

To investigate any possible degradation or metabolism of the ES8 compounds to release AA or Trp, we performed both short-term and long-term treatments of Col-0 and *wei2wei7* seedlings with the ES8 compounds, followed by compound analysis (Fig. S4). We measured the concentrations of the relevant ES8 compound, AA or Trp and the non-AA or non-Trp part of the ES8 compound *in planta* as well as in ES8 compound-supplemented treatment medium to which no seedlings were added. After short-term treatment (5 h incubation in liquid treatment medium) with 5 µM ES8, high levels of ES8 were detectable in the seedlings and the seedling-free treatment medium remained at about 5 µM ES8 (Fig. S4a). Importantly, these findings confirm that ES8 readily enters plant tissues during short-term treatment. After long-term treatment (9 d growth on solid treatment medium), the concentrations of ES8 in the seedlings and the seedling-free treatment medium had lowered considerably, suggesting degradation of ES8 over time, and/or slower uptake from solid than liquid treatment medium. Compared to ES8, much lower levels of ES8.7 and ES8.7-Trp were present in the seedlings after short-term treatment (Fig. S4b,c), suggesting that ES8 may be more efficiently taken up into seedling tissues or ES8.7 and ES8.7-Trp may be degraded or metabolized during the short-term treatment. Degradation of ES8.7-Trp was supported by our measurements of its concentration in the seedling-free treatment medium, which had already lowered to 3.3 µM after short-term incubation and to 0.5 µM after long-term incubation (Fig. S4c). Moreover, the levels of ES8.7-Trp were considerably lower in the seedlings after long-term compared to short-term treatment. While these results suggest that ES8 and ES8.7-Trp are likely degraded over time, the levels of AA and Trp in the seedlings after ES8 compound treatment were not different to the levels after mock treatment and neither AA nor Trp were detected in the treatment medium samples (Fig. S4d,e). Furthermore, we did not detect non-AA or non-Trp parts of the ES8 compounds at any time point, neither in the seedlings nor in the seedling-free treatment medium (Fig. S4f). Therefore, the observed activities of the ES8 compounds are not due to the release of AA or Trp leading to increased IAA biosynthesis.

### AA and ES8 can rescue root gravitropic growth in *wei2wei7* without rescuing IAA level

As AA is a precursor of IAA, we investigated the possibility that the rescue of root gravitropic growth by ES8 and AA might indirectly result from increased IAA biosynthesis. First, we measured IAA concentrations after long-term treatments with AA. In Col-0, only 10 µM AA significantly increased the IAA level (Fig. 2a), which likely explains why treatments of Col-0 with 10 µM and higher AA resulted in significantly shorter roots (Fig. S2a). While treatment of *wei2wei7* with 1 or 10 µM AA rescued the IAA level to that of mock-treated Col-0, treatment with 0.5 µM AA had no effect on IAA content (Fig. 2a), despite this concentration having almost fully rescued root gravitropic growth and partially rescued root length in *wei2wei7* seedlings (Fig. 1d). Next, we measured IAA content in seedlings treated long-term with 5 µM ES8, ES8.7 or ES8.7-Trp. While the IAA level was slightly but significantly reduced in mock-treated *wei2wei7* compared to Col-0, none of the ES8 compounds significantly affected IAA content compared to mock treatment in either genotype (Fig. 2b). As our IAA analysis was performed on whole seedlings, we cannot rule out small, local changes in IAA levels in specific regions of the root. However, taken together, our results suggest that ES8 and ES8.7 rescue *wei2wei7* root gravitropic growth without affecting general IAA content, thereby supporting the hypothesis that AA plays a role in the regulation of root gravitropic growth independently from its function in IAA biosynthesis and potentially via a previously unknown role in auxin distribution.

**Fig. 2.**
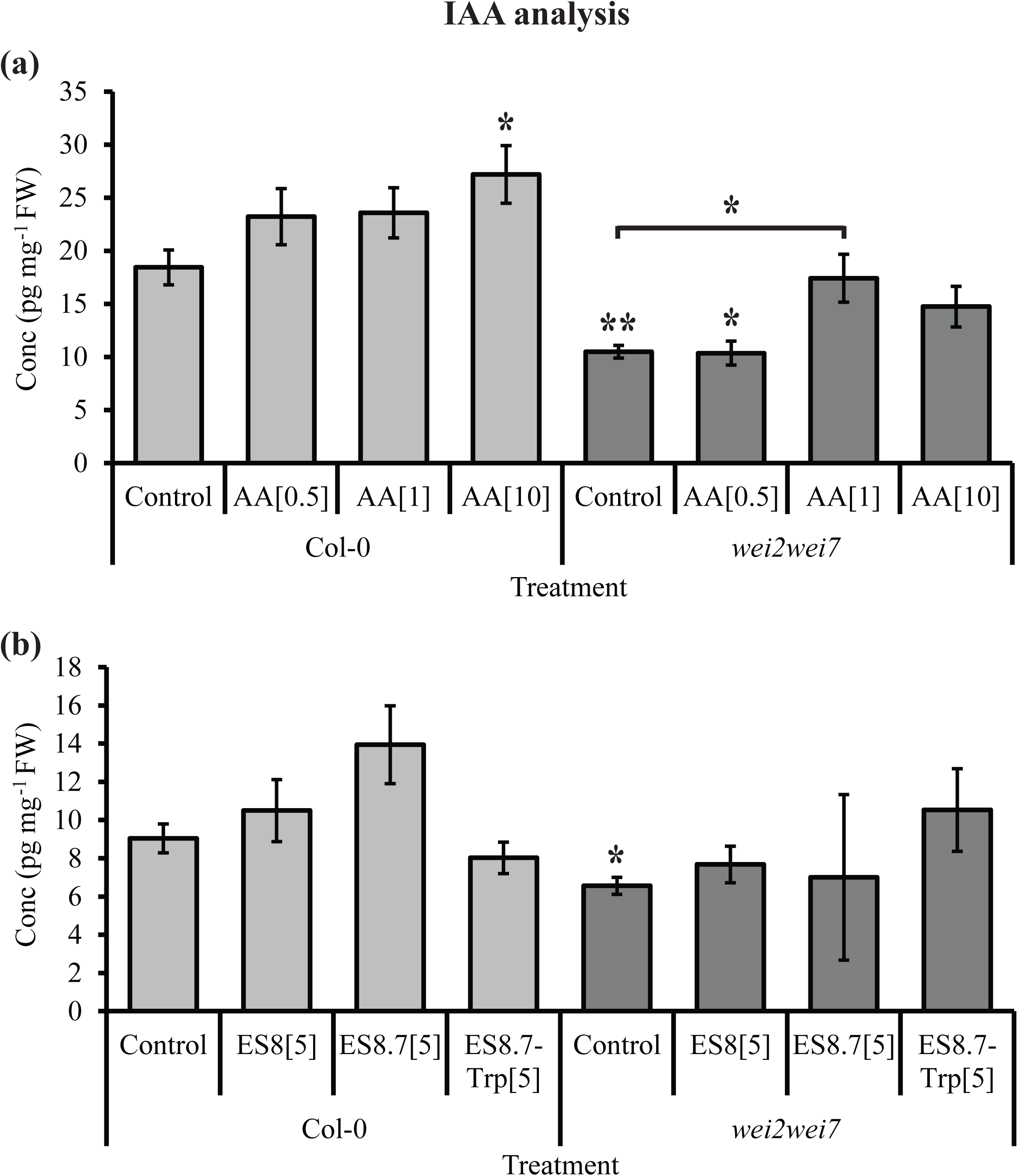
Treatment with anthranilic acid (AA), but not Endosidin 8 (ES8), can rescue the indole-3-acetic acid (IAA) level in *wei2wei7*. IAA concentrations (Conc) in 9-d-old seedlings of *Arabidopsis thaliana* Columbia-0 (Col-0) and *wei2wei7* grown on medium supplemented with AA (a) or with ES8, ES8.7 or ES8.7-Tryptophan (Trp) (b). Asterisks indicate samples significantly different from the Col-0 mock-treated control unless indicated otherwise (**, *P*<0.01; *, *P*<0.05). Error bars indicate standard error of the mean of the biological replicates. Values in square brackets indicate treatment concentrations in µM. Tissue was sampled from a mixture of 20 ground seedlings per sample per each of five (a) or four (b) biological replicates.

### Root gravitropic response is impaired by AA when its conversion to downstream IAA precursors is repressed

To test our hypothesis that AA may regulate root gravitropic growth via a role independent of IAA biosynthesis, we generated transformed *Arabidopsis* lines in which the gene encoding ASA1 (WEI2) (Niyogi & Fink, 1992) is constitutively overexpressed and that encoding PHOSPHORIBOSYLANTHRANILATE TRANSFERASE 1 (PAT1), which converts AA to the next downstream IAA precursor (Rose *et al*., 1992), is subject to estradiol-induced silencing. Of several homozygously transformed *35S::ASA1* (*35S::WEI2*) and *XVE::amiRNA-PAT1* lines, in which the promoters are known to be widely expressed (Odell *et al*., 1985; Zuo *et al*., 2000), we used qPCR analysis of *ASA1* (*WEI2*) and *PAT1* expression in whole seedlings (Fig. S5a,b) to select two lines for each construct displaying reproducible and strong constitutive *ASA1* induction (*35S::ASA1* lines 3B6 and 3B7) or inducible *PAT1* silencing (*XVE::amiRNA-PAT1* lines 2D4 and 4B10). We then crossed the selected lines and analyzed *ASA1* and *PAT1* expression in the progeny that were homozygous for both transformations, which we named *AxP* (*ASA1 x PAT1*) lines (Fig. S5c,d). While *ASA1* was overexpressed in all *AxP* lines, there was a tendency for increased *PAT1* expression in non-estradiol-induced conditions in those lines with highest *ASA1* expression, suggesting positive regulation between *ASA1* and *PAT1* genes. We selected two *AxP* lines for further experiments; *AxP1* (3B6 × 2D4 line no. 4) in which *ASA1* was five-fold overexpressed compared to the non-treated WT without affecting non-induced *PAT1* expression and *AxP2* (3B7 × 2D4 line no. 21), in which *ASA1* was 10-fold overexpressed, resulting in three-fold overexpression of *PAT1* in non-induced conditions (Fig. S5c,d). Additionally, an estradiol-inducible five- and three-fold reduction in *PAT1* expression compared to non-treated Col-0 was shown for *AxP1* and *AxP2*, respectively (Fig. S5d).

We analyzed the levels of IAA and several IAA precursors/conjugates/catabolites in WT and *AxP* lines treated long-term with estradiol (grown on supplemented medium) (Fig. S6a). Importantly, AA levels were significantly higher in both *AxP* lines compared to the WT, while the IAA content was not affected. The levels of Trp, IAN, IAM and oxIAA were also not significantly affected in the lines, while the levels of the IAA conjugates IAA-Aspartate (IAAsp) and IAA-Glutathione (IAGlu) showed rather variable results (Fig. S6a). These analyses suggest that simultaneous overexpression of *ASA1* and silencing of *PAT1* result in significantly increased AA levels, but do not alter IAA levels.

We next investigated the root phenotypes of the *AxP* lines. In control conditions, both lines displayed similar root gravitropic growth to, but slightly shorter roots than, the WT (Fig. S6b,c). After long-term estradiol treatment, the gravitropic growth of WT and *AxP* roots was slightly reduced, to a similar extent (Fig. S6b), while the root length of all genotypes was reduced, but more severely in *AxP* lines than the WT (Fig. S6c). To analyze root gravitropic responses in the *AxP* lines, we turned the seedlings 90° and subsequently measured the gravistimulated root bending angles (Fig. S6d). We divided the total number of roots, by percentage, into several categories of bending angles (Fig. 3). In control conditions, Col-0 and both *AxP* lines responded to the gravistimulus with a very similar range of root bending angles, with the majority of roots bending 75-105° (Fig. 3). Estradiol treatment inhibited the gravitropic response of Col-0 roots, reducing their bending angles, resulting in a significant reduction in the proportion of total roots bending 75-105° and a significant increase in the proportion bending <75° (Fig. 3a,b). The *AxP* lines, however, responded differently to estradiol than the WT. As for the WT, estradiol treatment resulted in both a significant reduction in the proportion of *AxP* roots bending 75-105° and, in the case of *AxP1*, a significant increase in the proportion bending <75°, but additionally resulted in a significant increase in the proportion bending >105° in both *AxP* lines (Fig. 3c-f). Therefore, while estradiol treatment specifically reduces root bending in the WT, this treatment results in both under- and over-bending in *AxP1* roots and over-bending in *AxP2* roots, in response to a gravistimulus. This suggests that increased AA levels in these lines interferes with root gravitropic responses, although we cannot rule out potential effects of changes in other IAA metabolite levels in root gravitropic responses.

**Fig. 3.**
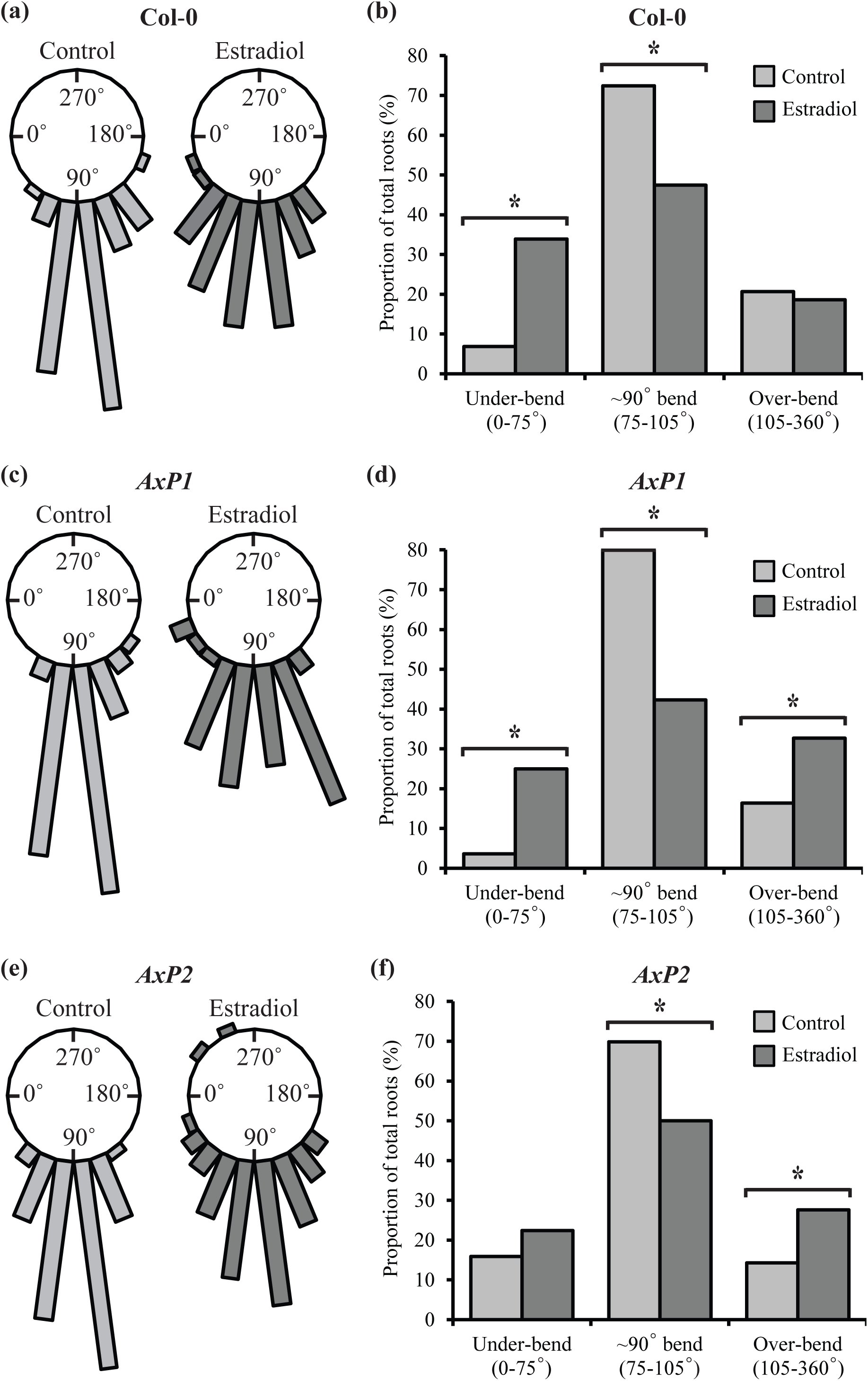
Anthranilic acid (AA) affects gravistimulated root bending independently of indole-3-acetic acid (IAA) biosynthesis. (a-f) Categories of root bending angles for gravistimulated control and 20 µM estradiol-treated seedlings of *Arabidopsis thaliana* Columbia-0 (Col-0) (a-b), *AxP1* (c-d) and *AxP2* (e-f). Polygonal x-axis frequency graphs showing root bending angles in 15° categories (a, c, e) and percentages of roots under-bending at 0-75° angles, bending at approximate right angles of 75-105° and over-bending at 105-360° (b, d, f). 5-d-old seedlings were transferred vertically to mock-(control) or 20 μM estradiol-supplemented medium for 24 h and then gravistimulated by turning 90° clockwise for a further 24 h before measuring gravistimulated root bending angles (see Fig. S6d). Frequencies/percentages were calculated based on the full dataset of all seedlings measured. Asterisks indicate significant differences between control and estradiol treatment, as calculated using the two-proportion z-test with significance set at *P<*0.1. *n* = 20 seedlings per sample per each of three biological replicates.

### PIN polarity in the stele is altered in *wei2wei7* and partially rescued by ES8

ES8 has been shown to disturb auxin distribution patterns in the root by altering PIN polarity (Doyle *et al*., 2015a). Considering that IAA itself can influence its own transport by regulating PIN abundance at the plasma membrane (Paciorek *et al*., 2005; Robert *et al*., 2010), we reasoned that AA, as a precursor of IAA, might also play such a role. To investigate this possibility, we first studied the effects of long-term ES8 and AA treatments on the expression pattern of the auxin-responsive promoter *DR5* in the root. To observe the effects of ES8 more easily, we used treatment at a high concentration of 15 µM, which led to a strong decrease in green fluorescent protein (GFP) signal in the stele of *DR5::GFP* WT roots (Fig. S7a), in agreement with previously published work (Doyle *et al*., 2015a). Furthermore, *DR5::GFP* crossed into the *wei2wei7* background showed a similarly low GFP signal in the stele in control conditions, which was reduced even further by 15 µM ES8 treatment (Fig. S7a). While 10 µM AA treatment did not noticeably affect the GFP signal in the stele of the WT, the signal in the *wei2wei7* stele was rescued by this treatment (Fig. S7a). These results suggest that AA may play a role in auxin distribution in the stele.

Next, we focused on the GFP signal in the root tip, particularly around the quiescent center (QC) and in the columella (Fig. S7b). We used both 5 and 15 µM ES8 treatment concentrations, which, in agreement with previously published work (Doyle *et al*., 2015a), led to an accumulation of GFP signal in cell file initials surrounding the QC in *DR5::GFP* WT, which were not labeled in control conditions (Fig. S7b). This accumulation of signal was rather striking, extending into lateral columella and root cap cells, at the higher ES8 treatment concentration of 15 µM. As found for the stele, *DR5::GFP* crossed into the *wei2wei7* background showed a similar GFP signal pattern in the root tip in control conditions as that induced by ES8 in the WT, with an accumulation of signal in the file initials surrounding the QC (Fig. S7b). This signal pattern was also apparent in the *wei2wei7* background after 5 µM ES8 treatment and was enhanced after 15 µM ES8 treatment. While 0.5 µM and 10 µM AA treatment did not noticeably affect the GFP signal in the root tip of Col-0, the signal in the *wei2wei7* root tip was slightly increased by 0.5 µM AA treatment and rescued to that of the control WT by 10 µM AA treatment (Fig. S7b). Therefore, we observed a negative correlation between the *DR5::GFP* signal strength in the root stele and tip. We suggest that a balanced AA level is important for proper auxin distribution in both the root stele and tip, as both addition of exogenous AA/ES8 and deficiency in endogenous AA levels disturbed *DR5::GFP* signal patterns. Together, these results indicate that AA may indeed play a role in auxin distribution in the root stele and tip, which likely affects gravitropic growth. However, it is important to note that we cannot rule out possible effects of AA/ES8 on local auxin biosynthesis within specific groups of cells.

Our observations of *DR5::GFP* signal in the stele prompted us to investigate the rootward-to-lateral plasma membrane fluorescence ratio (hereafter referred to as rootward polarity) of PIN1, PIN3 and PIN7 in the provascular cells of Col-0 and *wei2wei7* root tips. We treated seedlings short-term (2 h) with 15 µM ES8 or 10 µM AA, performed immunolabeling to observe endogenous PIN1 and PIN7 and used the *PIN3::PIN3-GFP* line crossed into the *wei2wei7* background due to poor labeling of antibodies against PIN3. The fluorescence signals for these PIN proteins were consistently weaker in the mutant than in the WT (Fig. 4a-c), suggesting decreased abundance at the plasma membranes. As previously reported by Doyle *et al*. (2015a), short-term ES8 treatment significantly, albeit slightly, reduced immunolocalized PIN1 rootward polarity in Col-0 and importantly, AA treatment produced a similar result (Fig. 4d). In contrast, PIN1 rootward polarity was significantly increased by about 20% in untreated *wei2wei7* compared to Col-0, while ES8 treatment appeared to rescue this hyper-polarity of PIN1 in the mutant back to almost that of the WT (Fig. 4d). Although PIN3-GFP rootward polarity was not affected by ES8 or AA treatments in either the Col-0 or *wei2wei7* backgrounds, it was increased by over 20% in the mutant compared to the WT (Fig. 4e). Finally, although PIN7 rootward polarity was not affected by ES8 or AA treatment in Col-0, it was strongly increased in the mutant compared to the WT, and like PIN1, was rescued in the mutant back to the level of the WT by ES8 treatment (Fig. 4f). These results suggest that AA may play a role in maintenance of PIN polarity in root provascular cells. One possible speculation on why treatment with AA, in contrast to ES8, did not rescue PIN1 or PIN7 polarity in the mutant may be a rapid conversion of AA to downstream IAA precursors within the seedlings.

**Fig. 4.**
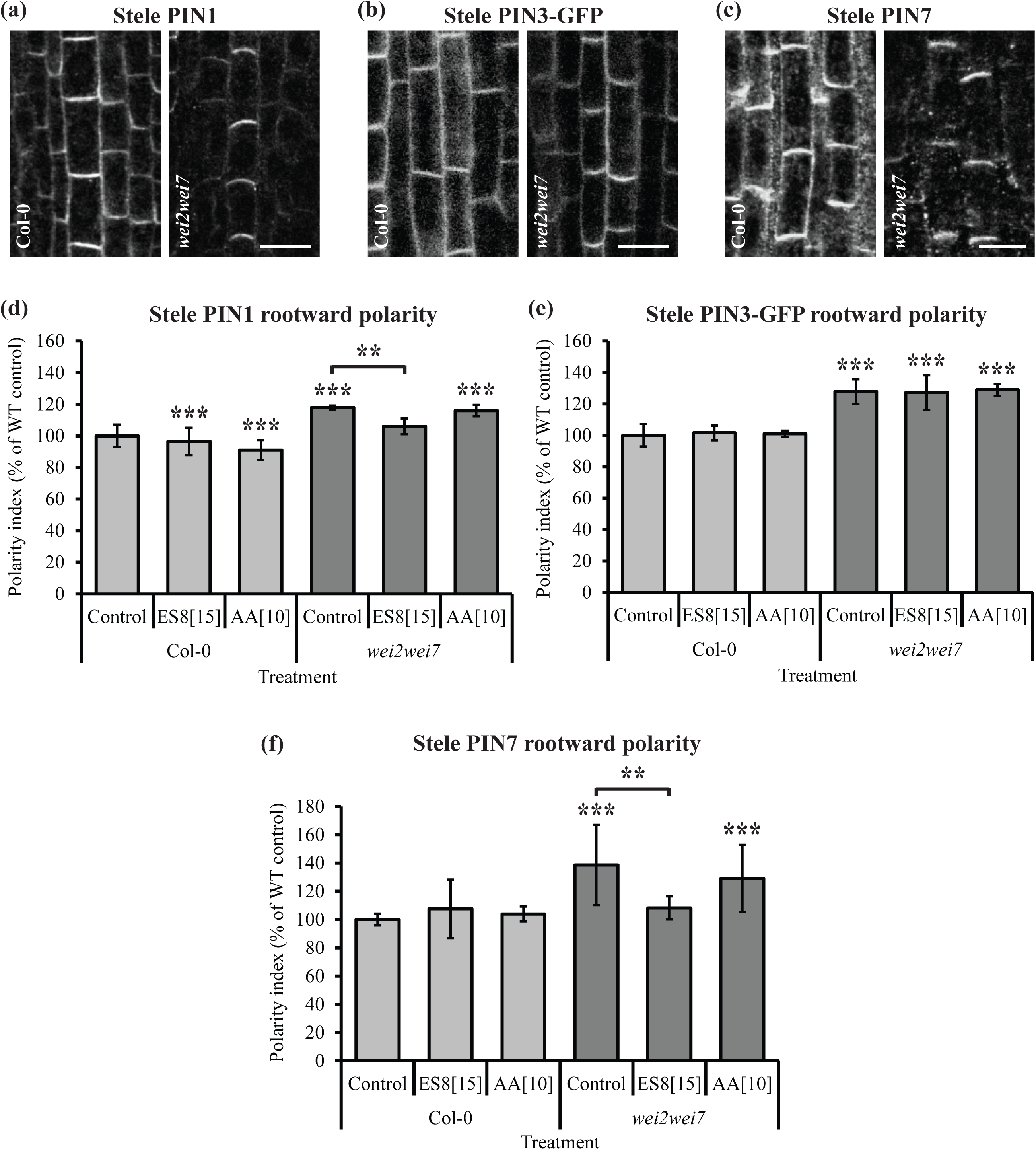
Anthranilic acid (AA) treatment or deficiency affects rootward polarity of PIN-FORMED (PIN) auxin transporters in root provascular cells. (a-c) Representative images of immunolabeled PIN1 (a), green fluorescent protein (GFP) fluorescence in *PIN3::PIN3-GFP* (b) and immunolabeled PIN7 (c) in stele provascular cells of 5-d-old *Arabidopsis thaliana* Columbia-0 (Col-0) and *wei2wei7* seedling roots. Scale bars represent 10 μm. (d-f) Rootward polarity index (expressed as a percentage of the Col-0 wild type (WT) mock-treated control) of fluorescence intensities of anti-PIN1 (d), GFP in *PIN3::PIN3-GFP* (e) and anti-PIN7 (f) in stele provascular cells of 5-d-old Col-0 and *wei2wei7* seedling roots treated for 2 h with Endosidin 8 (ES8) or AA in liquid treatment medium. Asterisks indicate samples significantly different from the Col-0 mock-treated control unless indicated otherwise (***, *P*<0.001; **, *P*<0.01). Error bars indicate standard error of the mean of the biological replicates. Values in square brackets indicate concentrations in µM. *n* = five cells per each of 15 seedlings per sample per each of three biological replicates.

As AA is a precursor of auxin, which is known to affect transcription of *PIN* genes (Vieten *et al*., 2005; Paponov *et al*., 2008), we investigated gene expression levels for all the plasma membrane-localized PIN proteins (*PIN1*, *PIN2*, *PIN3*, *PIN4* and *PIN7*) in WT and mutant seedlings at 9 d old, the age at which we performed our root gravitropic growth and length studies. The expression levels of *PIN1*, *PIN2* and *PIN4* were strongly decreased in *wei2wei7* compared to Col-0, while *PIN3* and *PIN7* expression levels were somewhat decreased, but not significantly (Fig. S8a). We next investigated the expression levels of *PIN1*, *PIN3* and *PIN7* in the same conditions used for our PIN polarity studies in root provascular cells (five-d-old seedlings treated with ES8 and AA for 2 h). At this stage, expression levels of *PIN1*, *PIN3* and *PIN7* were somewhat decreased in the mutant compared to the WT, but not significantly (Fig. S8b). Furthermore, treatment with ES8 and AA did not significantly affect the expression of these genes (Fig. S8b). These results imply that while transcription of *PIN* genes is decreased in *wei2wei7*, the effects of ES8 and AA on PIN polarity are not due to *PIN* gene transcriptional changes. Overall, our data suggest that endogenous AA may play a role in regulating the polarity of PIN1, PIN3 and PIN7 in root provascular cells *via* a mechanism unrelated to *PIN* gene expression levels. We previously determined that ES8 targets a secretory pathway delivering newly produced PIN1 toward the rootward plasma membranes of root provascular cells (Doyle *et al*., 2015a) and it is tempting to speculate that AA might play regulatory roles in similar PIN trafficking routes in order to guide auxin distribution in the root.

### AA regulates root gravitropism *via* repolarization of PIN3 and PIN7 in the columella

Our observations of *DR5::GFP* signal in the columella (Fig. S7b) indicate that AA may also play a role in auxin distribution specifically in this particular root tissue. Additionally, previous studies of the expression patterns of *ASA1* (*WEI2*) and *ASB1* (*WEI7*) promoter-GUS fusions in dark-grown *Arabidopsis* roots revealed strong expression in the root meristem and columella (Stepanova *et al*., 2005). Plasma membrane-localized PIN3 and PIN7 in the columella are thought to act in redistribution of auxin in response to gravistimulus (Friml *et al*., 2002b; Kleine-Vehn *et al*., 2010), potentially redundantly with PIN4, which is also localized in columella cells (Friml *et al*., 2002a; Vieten *et al*., 2005). We therefore reasoned that high expression of anthranilate synthase genes in the columella may reflect a role of AA in regulating gravity-responsive polarity of these PIN proteins. First, to investigate the expression patterns of the *ASA1* and *ASB1* promoters in light-grown roots, we performed GUS staining of *ASA1::GUS* (*WEI2::GUS*) and *ASB1::GUS* (*WEI7::GUS*) seedlings. We observed strong expression of the *ASB1* promoter, but not the *ASA1* promoter, in the stele of the upper root, while neither *ABA1* nor *ASB1* promoter expression was detected in the lower part of the root excluding the root tip (Fig. S9a,b). We observed strong *ASA1* and *ASB1* promoter expression in the tip of the root meristem and in the columella, with *ASA1::GUS* expressed throughout the columella, while *ASB1::GUS* expression was limited to the innermost columella cells (Fig. S9c).

Next, we investigated the localization of endogenous PIN3, PIN4 and PIN7 in the columella of Col-0 and *wei2wei7*. Interestingly, the fluorescence intensity of these proteins was consistently increased in the innermost cells of the columella in *wei2wei7* compared to Col-0 (Fig. S9d-f), suggesting that the abundance and/or localization of these proteins are altered in the mutant columella. The antibodies against these PIN proteins did not label the outermost columella cells, in agreement with previous studies using PIN3 and PIN4 antibodies (Friml *et al*., 2002a; Friml *et al*., 2002b). We therefore continued our studies using *PIN3::PIN3-GFP* and *PIN7::PIN7-GFP* lines crossed into the *wei2wei7* background (Fig. 5a,b). We performed long-term treatments of these lines with ES8 and AA and investigated the shootward-plus-rootward to lateral-plus-lateral fluorescence ratio (hereafter referred to as shootward-rootward polarity) of the GFP-labeled PIN proteins. While the shootward-rootward polarity of PIN3-GFP was similar in *wei2wei7* and Col-0 backgrounds regardless of compound treatment (Fig. 5c), PIN7-GFP was over 20% more polarly shootward-rootward localized in the mutant than in the WT (Fig. 5d). Moreover, 10 µM AA treatment partially rescued PIN7-GFP polarity in the mutant toward the WT level (Fig. 5d).

**Fig. 5.**
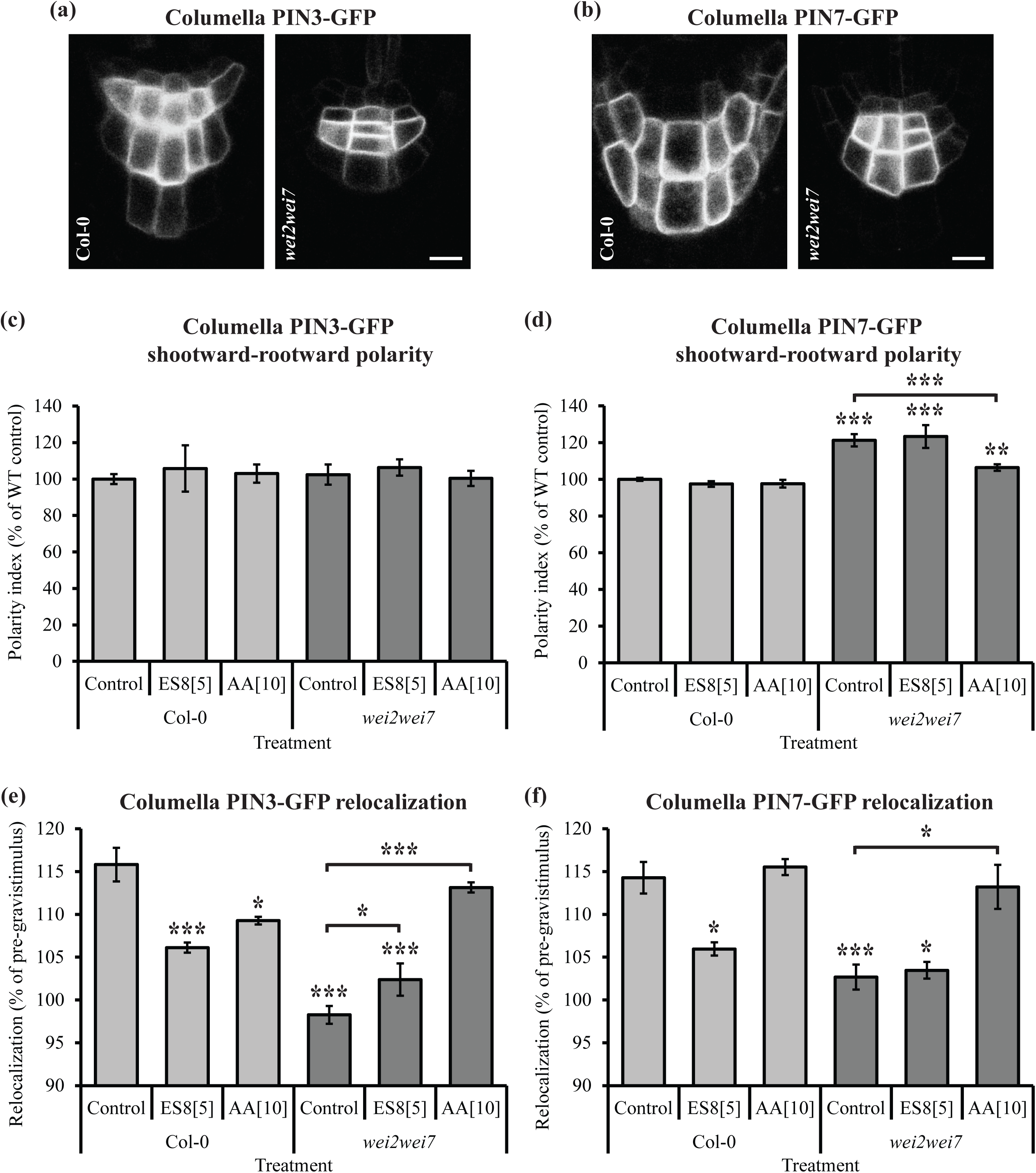
Anthranilic acid (AA) regulates gravistimulated relocalization of PIN-FORMED (PIN) auxin transporters in root columella cells. (a-b) Representative images of green fluorescent protein (GFP) fluorescence in *PIN3::PIN3-GFP* (a) and *PIN7::PIN7-GFP* (b) in columella cells of 5-d-old *Arabidopsis thaliana* Columbia-0 (Col-0) and *wei2wei7* seedling roots. Scale bars represent 10 µm. (c-d) Shootward-rootward polarity index (expressed as a percentage of the Col-0 wild type (WT) mock-treated control) of fluorescence intensities of GFP in *PIN3::PIN3-GFP* (c) and *PIN7::PIN7-GFP* (d) in columella cells of 5-d-old seedlings of Col-0 and *wei2wei7* grown on solid treatment medium supplemented with Endosidin 8 (ES8) or AA. (e-f) Fluorescence intensity relocalization (proportion of plasma membrane fluorescence on the lateral plasma membrane after gravistimulation expressed as a percentage of the same plasma membrane fluorescence proportion before gravistimulation) of GFP in *PIN3::PIN3-GFP* (e) and *PIN7::PIN7-GFP* (f) in columella cells of 5-d-old seedlings of Col-0 and *wei2wei7* grown on solid treatment medium and gravistimulated at 90° for 30 min. Lateral refers to the cellular position of the plasma membrane in vertically positioned roots, which became downward-facing during the gravistimulus (for more details see Methods S5 and Fig. S10). Asterisks indicate samples significantly different from the Col-0 mock-treated control unless indicated otherwise (***, *P*<0.001; **, *P*<0.01; *, *P*<0.05). Error bars indicate standard error of the mean of the biological replicates. Values in square brackets indicate concentrations in µM. *n* = five cells per each of 20 seedlings per sample per each of three biological replicates.

We next investigated gravity-induced relocalization of PIN3-GFP and PIN7-GFP in the columella. After a 90° gravistimulus for 30 min, about 15% more PIN3-GFP and PIN7-GFP were present on the now downward-facing (formerly lateral-facing) plasma membranes of the columella cells in WT seedlings (Fig. 5e,f and Fig. S10a,c). Long-term treatment of the WT with 5 µM ES8 or 10 µM AA strongly reduced PIN3-GFP relocalization to only about 5-10% (Fig. 5e and Fig. S10a). Strikingly, gravistimulus-induced relocalization of PIN3-GFP was completely absent in mock-treated *wei2wei7*, partially rescued by treatment with 5 µM ES8 and fully rescued by treatment with 10 µM AA (Fig 5e and Fig. S10b). Similar but less pronounced effects were observed for PIN7-GFP in the columella; relocalization was reduced by ES8 in the WT and almost absent in the mock-treated mutant (Fig. 5f and Fig. S10c,d). However, 10 µM AA did not affect PIN7-GFP relocalization in the WT and 5 µM ES8 did not rescue the PIN7-GFP relocalization defect in the mutant (Fig. 5f and Fig. S10c,d). The almost total absence of gravistimulus-induced PIN3- and PIN7-GFP relocalization in *wei2wei7* correlates with the mutant’s strong agravitropic root phenotype (Fig. 1c). Moreover, the partial rescue of gravistimulus-induced PIN3-GFP relocalization in *wei2wei7* by long-term treatment with 5 µM ES8 (Fig. 5e and Fig. S10b) appears to correlate with the partial rescue of root gravitropic growth by the same treatment (Fig. 1e). These results imply that endogenous AA may play a role in regulating relocalization of PIN3 and PIN7 proteins in the columella in response to gravity.

To further investigate a potential role for PIN proteins in AA-regulated root gravitropism, we analyzed root gravitropic growth in a range of *pin* mutants and their crosses with *wei2wei7*. Interestingly, while the *ethylene insensitive root1-4* (*eir1-4*, a *pin2* allele) mutant showed intermediate root gravitropic growth between *wei2wei7* and Col-0, crossing these mutants caused an additive effect, with *wei2wei7eir1-4* being more severely agravitropic than *wei2wei7* (Fig. S11a). Of the tested *pin3*, *pin4* and *pin7* alleles, none of the single mutants were affected in root gravitropic growth compared to the WT and introduction of the *pin3-4* or *pin7-2* mutations into the *wei2wei7* background did not affect the gravitropic growth. Interestingly, in contrast to the *eir1-4* mutation, introduction of the *pin3-5* or *pin4-3* mutations into *wei2wei7* partially rescued the root gravitropic growth compared to *wei2wei7* (Fig. S11a). While *pin1-501* showed increased root gravitropic growth compared to the WT, this mutation also partially rescued the root gravitropic growth of *wei2wei7* (Fig. S11a).

We next tested the effects of long-term treatments with high concentrations of AA on root gravitropic growth in the mutants. Similarly to the WT, none of the *pin* mutants tested showed any sensitivity to AA in terms of changes in root gravitropic growth (Fig. S11b). While the introduction of *pin3-4* or *pin7-2* to *wei2wei7* did not alter its sensitivity to AA in terms of increase in gravitropic index, crossing *eir1-4* into *wei2wei7* significantly increased its sensitivity to AA (Fig. S11b). In contrast, introduction of *pin3-5*, *pin4-3* or *pin1-501* to *wei2wei7* reduced its sensitivity to AA, resulting in decreased rescue of root gravitropic growth (Fig. S11b).

The recovery of, as well as the reduction in AA-induced rescue of, *wei2wei7* root gravitropic growth by introducing *pin1*, *pin3* or *pin4* mutations, provide further evidence for the involvement of PIN1 and PIN3, as well as implicating involvement of PIN4, in AA-regulated root gravitropism. It is unclear why *pin3-5* but not *pin3-4* partially rescues *wei2wei7* gravitropism and suppresses rescue of *wei2wei7* gravitropism by AA, but these different effects may be due to potential secondary mutations in one or both of the mutants, and/or the different positions of the T-DNA insertions. These insertions occur in an exon in *pin3-4* and in the UTR preceding the start codon in *pin3-5*. The well-known important role of PIN2 in root gravitropism (Abas *et al*., 2006; Kleine-Vehn *et al*., 2008), however, is most likely not related to AA-regulated root gravitropism, considering the strong additive effect of *eir1-4* and *wei2wei7* mutations in reducing root gravitropic growth and increasing sensitivity to AA.

Taken together, our results strongly support a new role for endogenous AA in root gravitropism via regulation of selective PIN protein polarity and dynamics and thereby auxin distribution in both the stele and columella and that this role of AA is independent of its well-known function in IAA biosynthesis.

## Discussion

We provide evidence in favor of a role for AA in root gravitropic growth through regulation of the subcellular localization of auxin transporter proteins, which likely influences the flow of auxin within the organ. Following their synthesis, most plasma membrane-targeted proteins are sorted and packaged into selective secretory trafficking routes (Gendre *et al*., 2014). It has been shown, for instance, that the auxin importer AUXIN-RESISTANT1 (AUX1) and exporter PIN1, when targeted to shootward or rootward plasma membranes of root tip cells respectively, are transported in distinct endosomes, subject to control by different regulatory proteins (Kleine-Vehn *et al*., 2006). The trafficking routes of such proteins may be distinct even if targeted to the same plasma membrane, as is the case, for example, for AUX1 and PIN3 in epidermal hypocotyl cells of the apical hook (Boutté *et al*., 2013). Such a remarkably complex system of endomembrane trafficking pathways is thought to allow for a high level of control, suggesting the likely existence of an array of selective endogenous compounds and/or signals regulating these trafficking routes.

Once polar plasma membrane-targeted auxin carriers have reached their destination, they remain remarkably dynamic, being subject to constant vesicular cycling (Geldner *et al*., 2001) to enable rapid retargeting in response to external stimuli (reviewed by Luschnig & Vert, 2014; and Naramoto, 2017). Auxin itself promotes its own flow by inhibiting clathrin-mediated endocytosis of PIN transporters, thus enhancing their presence at the plasma membrane (Paciorek *et al*., 2005). Our results suggest that AA, an important early precursor in IAA biosynthesis, may also act on PIN plasma membrane localization to regulate the flow of auxin, through currently unknown mechanisms.

The use of pharmacological inhibitors, identified through chemical biology approaches, has proven a powerful strategy that has greatly assisted in unravelling the details of auxin transporter trafficking mechanisms (reviewed by Hayashi & Overvoorde, 2014; and Doyle *et al*., 2015b). We previously employed such a strategy, revealing that the AA analog ES8 selectively inhibits an early endoplasmic reticulum (ER)-to-Golgi secretory pathway, regulated by the adenosine diphosphate (ADP) ribosylation factor guanine nucleotide exchange factors (ARF-GEFs) GNOM and GNOM-LIKE 1 (GNL1), involved in rootward targeting of PIN1 without affecting the polarity of shootward plasma membrane proteins (Doyle *et al*., 2015a). We suggest that AA itself is likely to act endogenously on PIN trafficking regulation in a similar way to ES8, but detailed studies on AA mechanisms may prove difficult due to the potential conversion of AA to other IAA precursors in plant tissues. As ES8 appears to mimic the effects of AA on PIN localization and root gravitropic growth without releasing AA through degradation and without affecting IAA levels, this synthetic compound provides great potential for understanding the mechanisms of AA on PIN localization in more detail, having already been extremely useful for distinguishing this newly discovered role of AA from its better-known role in auxin biosynthesis.

Any pharmacological treatments of biological tissues raise the question of compound uptake efficiency. Based on our analysis of the ES8 compounds inside plant tissues, we can conclude that uptake, either passive or active, of all these compounds occurs, with ES8 being taken up about 10 times faster than ES8.7 or ES8.7-Trp during short-term treatments. Although we currently do not know how these compounds enter plant tissues, one may speculate that AA and Trp transporters are likely to exist *in planta*, which might also transport ES8/ES8.7 and ES8.7-Trp, respectively. It will be of great interest in future studies to investigate the distribution and dynamics of compound uptake, which could potentially be observed by their labeling, fluorescently or otherwise.

Interestingly, the amino acid para-aminobenzoic acid, which has a similar structure to AA and is produced from the same precursor, has also recently been shown to play a role in root gravitropism, distinct from its better known role in folate biosynthesis (Nziengui *et al*., 2018). However, unlike AA, para-aminobenzoic acid promotes gravitropic root growth in WT plants as well as promoting gravistimulated root bending by enhancing the asymmetric auxin response between the two root sides (Nziengui *et al*., 2018). Another study linking AA, or more specifically Trp, with root bending, revealed that a mutation in the *ASA1* gene confers a more compressed wavy root phenotype than the WT when seedlings are grown on agar surfaces tilted 30° from the vertical (Rutherford *et al*., 1998). As the mutant roots respond normally to gravistimuli when grown in non-waving conditions (within agar), it could be concluded that the root waving phenotype is not caused by agravitropism. Furthermore, the phenotype is rescued in the mutant by supplementing with AA or Trp but not IAA. Mutations in the *TRYPTOPHAN SYNTHASE α* and *β1* genes confer similar phenotypes, suggesting that a Trp deficiency is responsible. While these results suggest that Trp may be involved in regulating non-agravitropic root waving, potentially related to thigmotropism, independently of IAA, our study did not use tilted plates and moreover, in our conditions (vertical plates), single mutants in *ASA1* and *ASB1*, namely *wei2-2* and *wei7-1*, did not show differences in root growth compared to the WT (Fig. S12). We would therefore argue that the reduced gravitropic index of *wei2wei7* roots is indeed due to a gravitropic defect, caused by AA deficiency. It is however very interesting that Trp, another IAA precursor, appears to regulate root directional growth in response to another stimulus. Indeed, the existence of several complex root growth regulatory mechanisms is hardly surprising considering the remarkable plasticity of this organ, the growth of which must respond to a wide array of internal signals and external stimuli, including gravity, touch, light, temperature, humidity and various chemical substances.

The agravitropic growth of *wei2wei7* roots may be due to a combination of decreased auxin content caused by reduced AA levels and the AA deficiency itself, as both auxin and AA affect the localization of PIN proteins. As was shown previously for ES8 (Doyle *et al*., 2015a), AA appears to act selectively depending on the PIN protein and the root tissue. PIN1, PIN3 and PIN7 all display increased rootward polarity in provascular cells of *wei2wei7* compared to Col-0, suggesting increased flow of auxin toward the root tip in the mutant. Correspondingly, we found decreased expression of the auxin-responsive promoter *DR5* in the root stele and increased expression around the root tip QC in the mutant, a pattern that was also observed in WT roots upon ES8 treatment. PIN7, but not PIN3, is also abnormally polarized in columella cells of *wei2wei7*, while both these proteins appear to be completely unresponsive to gravistimulus in the mutant columella. Furthermore, the high expression of *ASA1* (*WEI2*) and *ASB1* (*WEI7*) in the root columella of the WT suggests the importance of AA in this tissue in particular, which our results suggest is due to a role for this compound in gravity-regulated PIN distribution amongst the plasma membranes. The particular importance of PIN1 and PIN3 in AA-regulated root gravitropism was further supported by the rescue, as well as the reduction in AA-induced rescue, of *wei2wei7* root gravitropic growth by introduction of *pin1* or *pin3* mutations. Taken together, our results suggest that the endogenous compound AA plays a role in root gravitropism by regulating the polarity and gravity-induced relocalization of specific PIN proteins in the provascular and columella cells. Furthermore, this role of AA is distinct from its well-known function in auxin biosynthesis, which we suggest is more important for root elongation than gravitropic growth.

## Acknowledgments

We acknowledge the Knut and Alice Wallenberg Foundation (F.A.), in particular “ShapeSystems” grant number 2012.0050 (S.M.D., M.Karad., K.L. and S.R.), the Plant Fellows fellowship program (A.R.), the Swedish Research Council (SRC) (F.A.), in particular the SRC/Vinnova grants VR2013-4632 (M.M.) and VR2016-00768 (P.G.), the Kempe (P.G. and F.A.) and Carl Tryggers (P.G.) Foundations, the Ghent University Special Research Fund (M.Karam.), the Czech Science Foundation project no. 13-40637S (M.Z.), the Göran Gustafsson Foundation and the Swedish Foundation for Strategic Research (F.A.) for funding. The core facility CELLIM of CEITEC was supported by the MEYS CR (LM2015062 Czech-BioImaging). We are grateful to Vanessa Schmidt and Roger Granbom for technical assistance, Christian Luschnig and Jiří Friml for sharing antibodies and seeds, Per-Anders Enquist for technical advice and especially Hélène S. Robert for sharing antibodies and primer sequences, helpful advice and critical reading of the manuscript.

## Author contributions

S.M.D., A.R. and S.R. designed the research; S.M.D., A.R., P.G., M.Karad., D.K.B., M.M., B.P., M.Karam., M.Z. and A.P. performed the research under the supervision of F.A., K.L., O.N. and S.R.; S.M.D., A.R., P.G., O.N. and S.R. interpreted the data; S.M.D. wrote the manuscript with input from A.R., P.G. and S.R.; all authors gave feedback on the final manuscript version; S.M.D. and A.R. contributed equally; P.G. and M.Karad. contributed equally.

## Supporting Information

**Methods S1** Selection of homozygous crossed mutants, generation of *35S::ASA1* (*35S::WEI2*) and *XVE::amiRNA-PAT1* lines and root growth measurements. For selection of homozygous lines at the F2 generation, seedlings displaying a *wei2wei7* phenotype were initially selected and then genotyped for the third mutation for triple mutants (see Table S2 for primers used), or observed on a fluorescence stereomicroscope for GFP marker line crosses. Finally, selected lines were confirmed for homozygous *wei2-1* and *wei7-1* mutations by genotyping (see Table S2 for primers used). The heterozygous line *pin1-501* (we added *01* to the name of this mutant to distinguish from another *pin1-5* allele) and its cross with *wei2wei7* were transferred to soil after imaging on treatment plates and only those seedlings that later formed pin-like inflorescences were included in root measurements in the initial images.

For generation of *35S::ASA1* and *XVE::amiRNA-PAT1* lines, see Table S2 for primers used. For constitutive overexpression of *ASA1* (AT5G05730), the coding region of the gene was amplified from *Arabidopsis thaliana* Col-0 cDNA. Primers for artificial microRNA (amiRNA) to knock down the *PAT1* (AT5G17990) gene were designed using the Web MicroRNA Designer tool (http://wmd3.weigelworld.org). The amiRNA was obtained using the pRS300 vector as a PCR template, as described previously (Ossowski *et al*., 2008). The fragments were introduced into the Gateway pENTR/D-TOPO cloning vector (Invitrogen, Thermo Fisher Scientific, Waltham, MA, USA) and verified by sequencing. The *ASA1* coding sequence was then cloned into the DL-phosphinothricin-resistant vector pFAST-R02 (Shimada *et al*., 2010), while the amiRNA was cloned into the hygromycin-resistant vector pMDC7b containing the estradiol-inducible XVE system (Curtis & Grossniklaus, 2003), using the LR reaction (Invitrogen). *Agrobacterium*-mediated transformation of the constructs into *Arabidopsis thaliana* Col-0 was achieved by floral dipping (Clough & Bent, 1998). Transformed plants were selected *via* antibiotic resistance on agar plates supplied with the respective antibiotics, 50 µg ml^−1^ hygromycin B or 25 µg ml^−1^ DL-phosphinothricin. Four and six independent homozygous *35S::ASA1* and *XVE::amiRNA-PAT1* lines were analyzed, respectively. Two lines per construct were then selected that displayed strong and reproducible constitutive *ASA1* induction or induced *PAT1* silencing, as determined by performing qPCR (Methods S4) using RNA extracted from 1-wk-old whole seedlings (Fig. S6a,b). To induce silencing in the *XVE::amiRNA-PAT1* seedlings, the seedlings were germinated and grown on agar plates supplemented with 20 µM estradiol, with DMSO used as a mock treatment control. Each selected *35S::ASA1* line was crossed with each selected *XVE::amiRNA-PAT1* line and homozygous F2 generation offspring, named *AxP* (*ASA1 x PAT1*) lines, were selected as before *via* antibiotic resistance. The homozygous lines were then tested for gene expression as before *via* qPCR (12 independent lines were analyzed) (Fig. S6c,d). Finally, two lines displaying reproducible simultaneous constitutive *ASA1* induction and induced *PAT1* silencing were selected for use in further experiments.

All root length and gravitropic index measurements were performed on 9-d-old seedlings, while microscopy studies were performed on 5-d-old seedlings due to cell collapse in a proportion of the tips of older *wei2wei7* roots. ImageJ software (https://imagej.nih.gov/ij/) was used to measure root length and vertical gravitropic index, which was calculated for each root as a ratio of L_y_: L, where L_y_ is the vertical distance from root base to tip, or the real depth of root tip penetration, and L is the root length, as described in Grabov *et al*. (2005). For gravistimulated root bending experiments, seedlings were grown vertically for 5 d in treatment-free conditions, then transferred to mock or estradiol-supplemented medium for 24 h, and then gravistimulated by turning the plates 90° for 24 h before measuring the root bending angles. The angles were measured in the direction of root bending between two lines intersecting at the former root tip position before gravistimulus, one being horizontal and the other originating from the current root tip position (Fig. S6d).

**Methods S2** Chemical synthesis and IAA metabolite analysis. The following procedure was used to synthesize (4-(*N*-(4-chlorobenzyl) methylsulfonamido)benzoyl)tryptophan (ES8.7-Trp). Methyl sulfonyl chloride (3.1 ml, 45.5 mmol, 1.2 equivalent) and pyridine (3 ml, 37.1 mmol, 1.1 equivalent) were added to a solution of 4-aminoethylbenzoate (5.1 g, 33.7 mmol) in acetonitrile (100 ml) at 0 °C and then stirred overnight at room temperature. The reaction mixture was concentrated and the resulting residue was dissolved in ethyl acetate. The organic layer was washed with HCl (2 *N*, 100 ml), saturated NaHCO_3_, H_2_O and brine, dried over Na_2_SO_4_, and concentrated to give an off-white solid (7.1 g, 94%), ethyl 4-(methylsulfonamido) benzoic acid, which was used without further purification. Then 4-chlorobenzyl bromide (1.43 ml, 11 mmol, 1.2 equivalent) and K_2_CO_3_ (3.8 g, 27.4 mmol, 3 equivalent) were added to a solution of ethyl 4-(methylsulfonamido)benzoate (2.1 g, 9.15 mmol) in dimethylformamide (DMF) (14 ml). The reaction mixture was stirred overnight after which liquid chromatography-mass spectrometry indicated the reaction was complete, yielding 4-(*N*-(4-chlorobenzyl)methylsulfonamido) ethylbenzoate (2.8 g, 88%). Next, NaOH (2 *N*, 15 ml) and H_2_O (10 ml) were added to 2 g of 4-(*N*-(4-chlorobenzyl)methylsulfonamido)ethylbenzoate, followed by overnight stirring. The reaction mixture was acidified to pH 5 with concentrated HCl and partitioned between ethyl acetate and H_2_O. The organic layer was washed with brine, dried over Na_2_SO_4_, and concentrated to give 4-(*N*-(4-chlorobenzyl)methylsulfonamido)ethylbenzoate (1.7 g, 92%), which was used without further purification. Then 10 ml thionyl chloride was added to a solution of 1 mmol 4-(*N*-(4-chlorobenzyl)methylsulfonamido)ethylbenzoate and refluxed for 12 h under a nitrogen atmosphere. The thionyl chloride was removed under reduced pressure to give acid chloride. Then 1.2 mmol tryptophan was dissolved in 5 ml DMF and cooled to 0 °C. At this temperature the acid chloride was added slowly by dissolving in 3 ml DMF. The reaction mixture was stirred at room temperature for 24 h. Ice cold water was added to the reaction mixture, which was then extracted with chloroform and washed with brine solution three times to afford the crude mixture. Finally, purification with column chromatography using methanol and chloroform as a solvent system resulted in a light yellow solid, (4-(*N*-(4-chlorobenzyl) methylsulfonamido)benzoyl)tryptophan (ES8.7-Trp) with 68% yield.

For IAA metabolite analysis, frozen samples were homogenized using a MixerMill bead mill (Retsch GmbH, Haan, Germany) and extracted in 1 ml of 50 mM sodium phosphate buffer containing 1% sodium diethyldithiocarbamate and a mixture of ^13^C_6_- or deuterium-labeled internal standards. After pH adjustment to 2.7 by 1 M HCl, a solid-phase extraction was performed using Oasis HLB columns (30 mg 1 cc; Waters Corporation, Milford, MA, USA). Mass spectrometry analysis and quantification were performed by a liquid chromatography–tandem mass spectrometry (LC-MS/MS) system comprising of a 1290 Infinity Binary LC System coupled to a 6490 Triple Quad LC/MS System with Jet Stream and Dual Ion Funnel technologies (Agilent, Santa Clara, CA, USA).

**Methods S3** Compound degradation analysis. For short-term treatments with 5 µM ES8 compounds, 5-d-old Col-0 and *wei2wei7* seedlings were incubated for 5 h in ES8, ES8.7 and ES8.7-Trp-supplemented liquid medium before harvesting in liquid nitrogen along with samples of liquid treatment medium incubated for 5 h without any seedlings. For long-term treatments with 5 µM ES8 compounds, Col-0 and *wei2wei7* seedlings were grown for 9 d on ES8, ES8.7 and ES8.7-Trp-supplemented solid medium before harvesting in liquid nitrogen along with samples of solid treatment medium incubated for 9 d without any seedlings. Samples from two biological replicates were harvested and divided into two technical replicates each. The medium samples were diluted 1:100 with 30% v/v acetonitrile and 10 µl was injected onto a Kinetex 1.7 μm C18 100A reversed-phase column 50 × 2.1 mm (Phenomenex, Torrance, CA, USA) followed by analysis by LC-MS/MS. The seedling samples (around 10 mg fresh weight) were extracted in 0.4 ml acetonitrile using a MixerMill MM 301 bead mill (Retsch GmbH) with 2 mm ceria-stabilized zirconium oxide beads at a frequency of 27 Hz for 3 min. The plant tissue extracts were then incubated at 4 °C with continuous shaking for 30 min, centrifuged in a Beckman Coulter (Brea, CA, USA) Avanti 30 at 4 °C for 15 min at 36,670 *g* and purified by liquid-liquid extraction using acetonitrile:hexan:H_2_O:formic acid (4:5:1:0.01) to remove impurities and the sample matrix. After 30 min incubation at 4 °C, the acetonitrile fractions were removed, evaporated to dryness *in vacuo* and dissolved in 50 μl 30% v/v acetonitrile prior to LC-MS/MS analysis using an Acquity UPLC I-Class System (Waters Corporation) coupled to a Xevo TQ-S MS triple quadrupole mass spectrometer (Waters Corporation). After injection of 10 µl, the purified samples were eluted using an 11 min gradient comprised of 0.1% acetic acid in methanol (A) and 0.1% acetic acid in water (B) at a flow rate of 0.5 ml/min and column temperature of 40 °C. The binary linear gradient of 0 min 2:98 A:B, 11 min 95:5 A:B was used, after which the column was washed with 100% methanol for 1 min and re-equilibrated to initial conditions for 2 min. The effluent was introduced into the MS system with the following optimal settings: source/desolvation temperature 150/600 °C, cone/desolvation gas flow 150/1000 l h^−1^, capillary voltage 1 kV, cone voltage 25-40 V, collision energy 15-30 eV and collision gas flow (argon) 0.21 ml/min. Quantification and confirmation of the ES8 compounds were obtained by the multiple reaction monitoring mode using the following mass transitions: 274>111/274>230, 457>378/457>413 and 526>125/526>322 for ES8, ES8.7 and ES8.7-Trp, respectively. The non-AA and non-Trp parts of the ES8 compounds were monitored by high resolution mass spectrometry (HRMS) using a Synapt G2-Si hybrid Q-TOF tandem mass spectrometer (Waters Corporation) equipped with electrospray ionization (ESI) interface (source temperature 150 °C, desolvation temperature 550 °C, capillary voltage 1 kV and cone voltage 25 V). Nitrogen was used as the cone gas (50 l h^−1^) and the desolvation gas (1000 l h^−1^). Data acquisition was performed in full-scan mode (50-1000 Da) with a scan time of 0.5 sec and collision energy of 4 eV; argon was used as the collision gas (optimized pressure of 5 × 10^−3^ mbar). The HRMS analyses were performed in positive (ESI^+^) and negative (ESI^−^) modes. Finally, analyses of AA and Trp were performed according to Novák *et al*. (2012) (Methods S2). All chromatograms were analyzed with MassLynx software (version 4.1; Waters Corporation) and the compounds were quantified according to their recovery.

**Methods S4** qPCR, GUS staining and generation of PIN7 antibody. For qPCR, total RNA was extracted from whole seedlings using the RNeasy Plant Mini Kit (Qiagen, Germantown, MD, USA) according to the manufacturer’s instructions. Samples were harvested in liquid nitrogen and the frozen tissue ground directly in their microtubes using microtube pestles. RQ1 RNase-free DNase (Promega Corporation, Madison, WI, USA) was used for the on-column DNase digestion step. RNA concentration was measured with a NanoDrop 2000 spectrophotometer (Thermo Fisher Scientific, Waltham, MA, USA). cDNA was prepared with 1 µg RNA using the iScript cDNA Synthesis Kit (Bio-Rad, Hercules, CA, USA) according to the manufacturer’s instructions. Serial dilutions of pooled cDNA from all samples for a particular experiment were used to determine efficiencies for each primer pair. qPCR was performed on a LightCycler 480 System (Roche Diagnostics, Rotkreuz, Switzerland) using LightCycler 480 SYBR Green I Master reagents (Roche Diagnostics), including two technical replicates per sample. For amplification of mRNA, the following protocol was applied: 95 °C for 5 min, then 40 cycles of 95 °C for 10 sec, 60 °C for 15 sec and 72 °C for 20 sec. For each experiment, transcriptional levels of the four reference genes *PEX4* (AT5G25760), *GADPH* (AT1G13440), *TIP41-LIKE* (AT4G34270) and *PP2A PDF2* (AT1G13320) were analyzed alongside the target genes (see Table S2 for primers used). Expression levels of the target genes were normalized against the two most stably expressed reference genes, as determined using GeNorm (Biogazelle, Ghent, Belgium) (Vandesompele *et al*., 2002), using the formula below, where E = efficiency, R = reference gene, Cq = quantification cycle mean, T = target gene. For each target gene, the normalized expression values were scaled relative to that of the WT control.

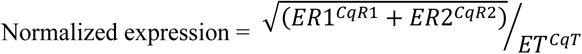

For GUS staining, seedlings were fixed in 80% acetone at −20 °C for 20 min, washed three times with distilled water and then incubated in 2 mM X-GlcA in GUS buffer (0.1% triton X100, 10 mM EDTA, 0.5 mM potassium ferrocyanide and 0.5 mM potassium ferricyanide in 0.1 M phosphate buffer (Na2HPO4 / NaH2PO4) at pH 7). The samples were then infiltrated for 10 min in a vacuum desiccator before incubation in the dark at 37 °C. The GUS reaction was stopped by replacing the GUS buffer with 70% ethanol. Samples were then mounted in 50% glycerol and observed on a Zeiss (Oberkochen, Germany) Axioplan microscope.

For the generation of anti-PIN7, a region of 882 bp corresponding to the hydrophilic loop of PIN7 was amplified and attB1 and attB2 recombination sites were incorporated (see Table S2 for primer sequences). The amplicon was recombined into the pDONR221 vector (Invitrogen) and the resulting pDONR221::PIN7HL was recombined into the pDEST17 vector (Invitrogen) in order to express the PIN7HL (31.4 kDa) in BL21 DE Star A *E. coli* cells. Cell cultures (250 ml) were induced in the logarithmic stage (after approx. 3.5 h) with 0.5 mM isopropyl β-D-1-thiogalactopyranoside (IPTG) for 7 h. Cells were harvested by centrifugation and resuspended in 15 ml phosphate buffered saline (PBS) at pH 8.0 with 8 M urea and 10 mM imidazole and incubated at 4 °C for 2 d. The PIN7HL expressed peptide was purified according to the Ni-NTA Purification System (Qiagen). The purified protein was then diluted in PBS buffer at pH 8.0 and desalted using Pierce (Thermo Fisher Scientific) concentrators 9K MWCO. The concentrated peptide was once more diluted in PBS at pH 8.0 before antibody production in rabbit, which was performed by the Moravian Biotechnology (Brno, Czech Republic) company (http://www.moravian-biotech.com/). Finally, serum specificity tests were performed in Col-0, *pin7* mutants and *PIN7-GFP* lines.

**Methods S5** Microscopy image quantifications. For live imaging, seedlings were mounted in their treatment medium for microscopic observations and images were acquired using identical acquisition parameters between mock and chemical treatments and between the WT and mutant in each experiment. For PIN rootward (basal) polarity index quantification in stele provascular cells, the ‘mean gray area’ tool in ImageJ was used to measure plasma membrane fluorescence intensity in confocal images and a rootward to lateral fluorescence ratio was calculated for each cell measured. For PIN shootward-rootward polarity index measurements in columella cells, the shootward-plus-rootward to lateral-plus-lateral plasma membrane fluorescence ratio was calculated. To monitor gravitropically induced PIN relocalization in columella cells, confocal z-stacks were acquired directly before and after a 30 min gravistimulation, during which the seedlings were rotated 90° to the horizontal position. Fluorescence intensity at the shootward (apical), rootward and lateral plasma membranes of the left- and right-most columella cells was measured on maximal intensity projections of the z-stacks using ImageJ (see Fig. S10 for examples of the cells measured). The percentage of fluorescence intensity distributed on each of the four plasma membranes of the same cell was calculated before and after gravistimulation. Relocalization was calculated by comparing the percentage of signal on the right-most lateral plasma membrane of the cell (which became the downward-facing plasma membrane during the gravistimulus) before and after gravistimulation.

**Table S1:**
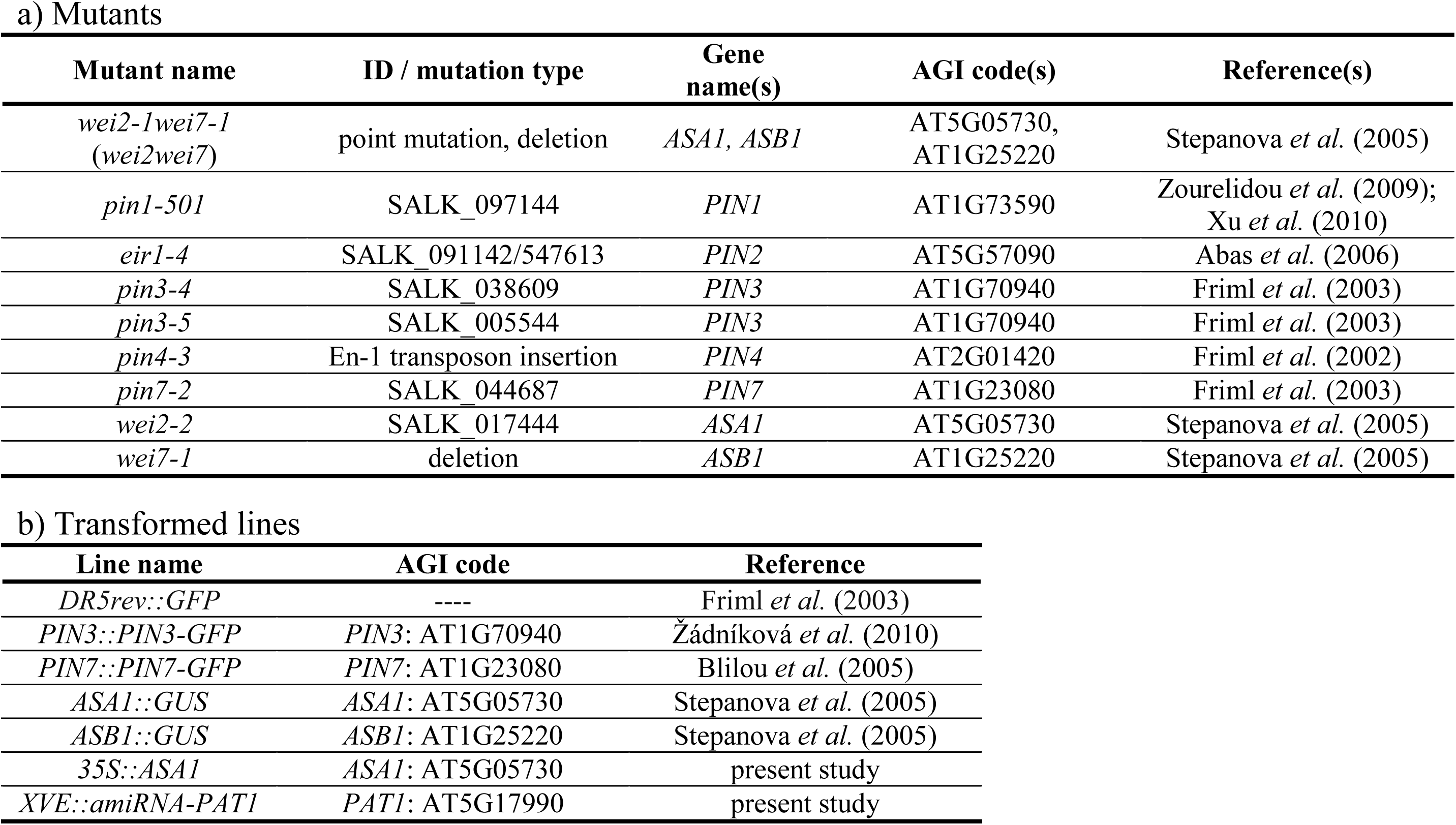
*Arabidopsis* mutants (a) and transformed lines (b) used in this study. SALK/SAIL T-DNA insertion lines (Alonso *et al*., 2003) were obtained from the Nottingham Arabidopsis Stock Centre (NASC). Arabidopsis Genome Initiative (AGI) codes are included for all genes. We renamed *pin1-5* SALK line 097144 to *pin1-501* to distinguish the mutant from another published *pin1-5* allele.

| Mutant name | ID / mutation type | Gene name(s) | AGI code(s) | Reference(s) |
| --- | --- | --- | --- | --- |
| *wei2-1wei7-1*<br>(*wei2wei7*) | point mutation, deletion | *ASA1*, *ASB1* | AT5G05730,<br>AT1G25220 | Stepanova *et al.* (2005) |
| *pin1-501* | SALK_097144 | *PIN1* | AT1G73590 | Zourelidou *et al.* (2009);<br>Xu *et al.* (2010) |
| *eir1-4* | SALK_091142/547613 | *PIN2* | AT5G57090 | Abas *et al.* (2006) |
| *pin3-4* | SALK_038609 | *PIN3* | AT1G70940 | Friml *et al.* (2003) |
| *pin3-5* | SALK_005544 | *PIN3* | AT1G70940 | Friml *et al.* (2003) |
| *pin4-3* | En-1 transposon insertion | *PIN4* | AT2G01420 | Friml *et al.* (2002) |
| *pin7-2* | SALK_044687 | *PIN7* | AT1G23080 | Friml *et al.* (2003) |
| *wei2-2* | SALK_017444 | *ASA1* | AT5G05730 | Stepanova *et al.* (2005) |
| *wei7-1* | deletion | *ASB1* | AT1G25220 | Stepanova *et al.* (2005) |

b) Transformed lines
| Line name | AGI code | Reference |
| --- | --- | --- |
| *DR5rev::GFP* | ---- | Friml *et al.* (2003) |
| *PIN3::PIN3-GFP* | *PIN3*: AT1G70940 | Žádníková *et al.* (2010) |
| *PIN7::PIN7-GFP* | *PIN7*: AT1G23080 | Blilou *et al.* (2005) |
| *ASA1::GUS* | *ASA1*: AT5G05730 | Stepanova *et al.* (2005) |
| *ASB1::GUS* | *ASB1*: AT1G25220 | Stepanova *et al.* (2005) |
| *35S::ASA1* | *ASA1*: AT5G05730 | present study |
| *XVE::amiRNA-PAT1* | *PAT1*: AT5G17990 | present study |

**Table S2:**
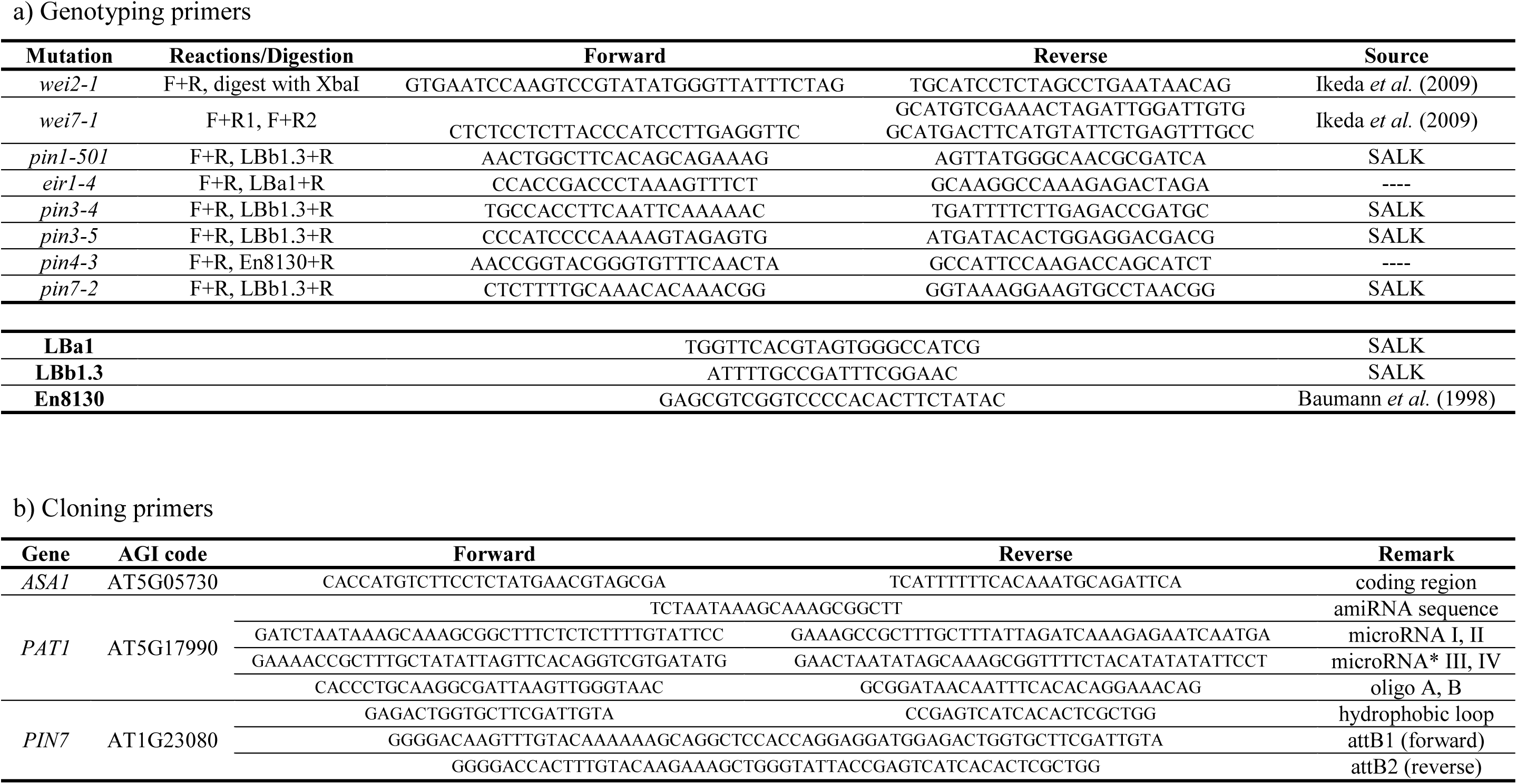
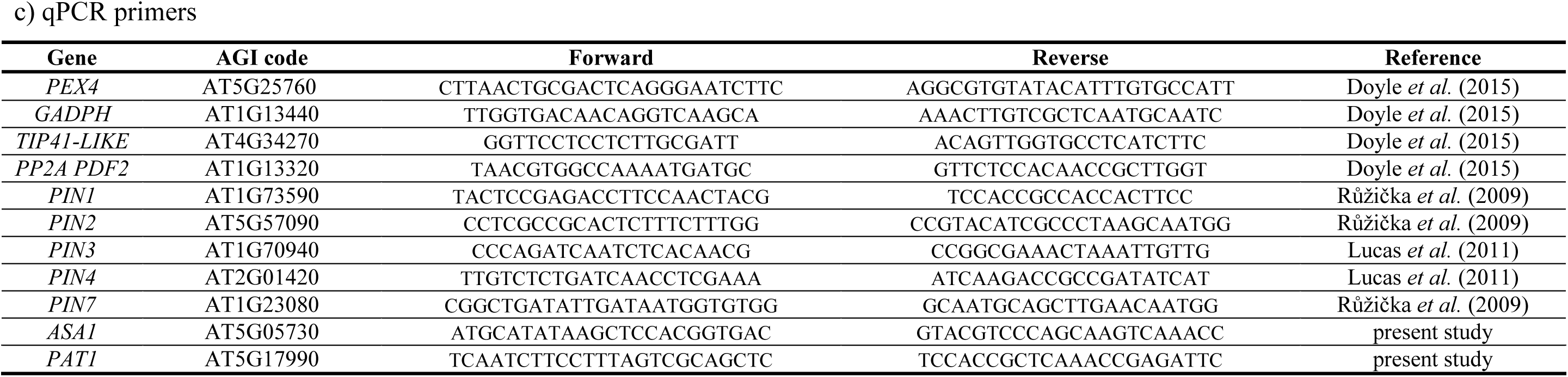
Genotyping (a), cloning (b) and qPCR (c) primers used in this study. Primers marked SALK were sourced from the SALK Institute Genomic Analysis Laboratory website (http://signal.salk.edu/). Primers with no indicated source have not been published previously (sequences designed in the present study or kindly provided by collaborators). Arabidopsis Genome Initiative (AGI) codes are included (for affected gene AGIs in mutants listed in (a) see Table S1).

**Fig. S1.**
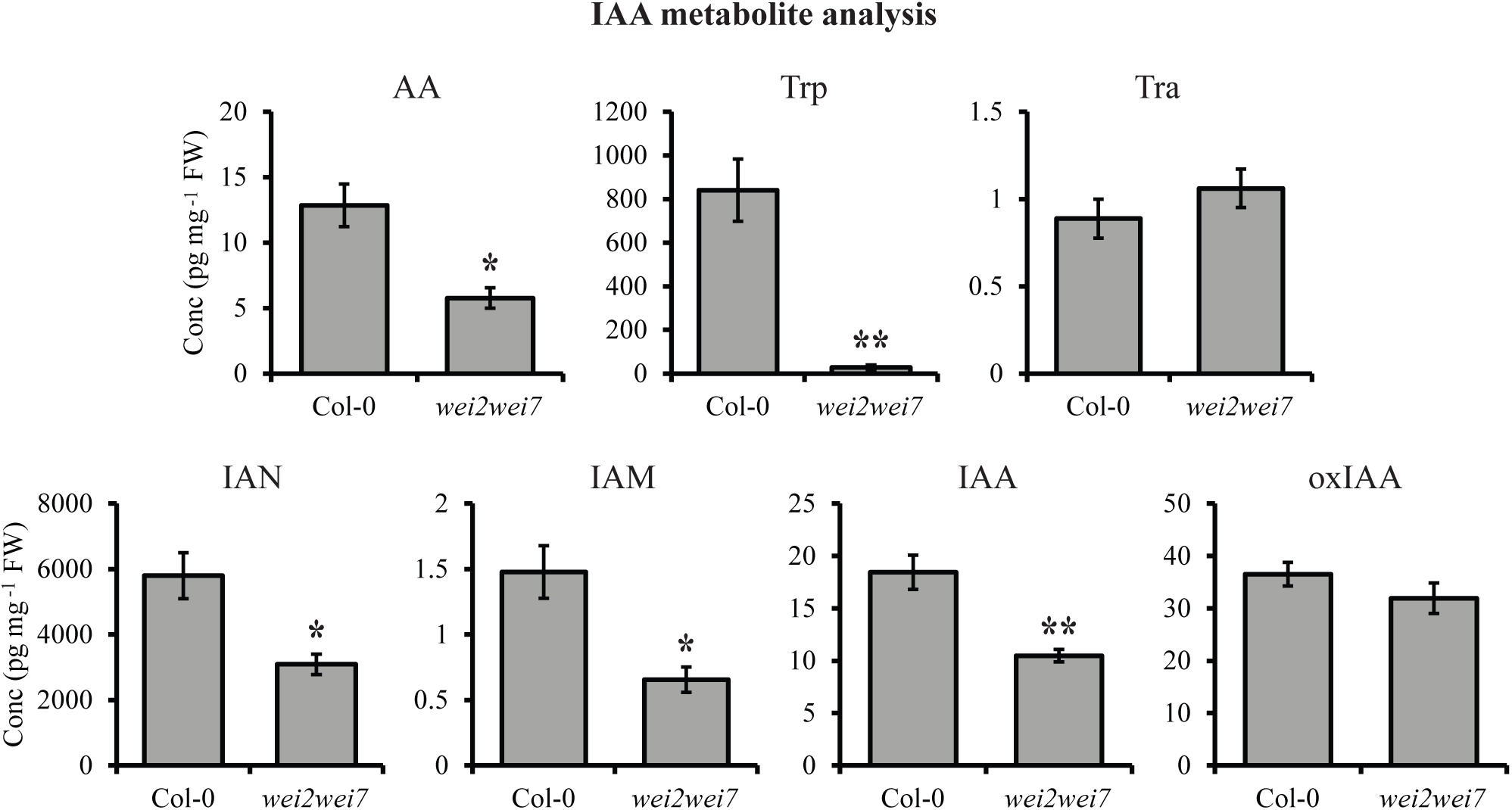
AA and other IAA precursors are deficient in *wei2wei7*. Concentrations (Conc) of anthranilic acid (AA), tryptophan (Trp), tryptamine (Tra), indole-3-acetonitrile (IAN), indole-3-acetamide (IAM), indole-3-acetic acid (IAA) and 2-oxoindole-3-acetic acid (oxIAA) in 9-d-old seedlings of *Arabidopsis thaliana* Col-0 and *wei2wei7*. Asterisks indicate significant differences from Col-0 (**, *P*<0.01; *, *P*<0.05). Error bars indicate standard error of the mean of the biological replicates. Tissue was sampled from a mixture of 20 ground seedlings per sample per each of four biological replicates.

**Fig. S2.**
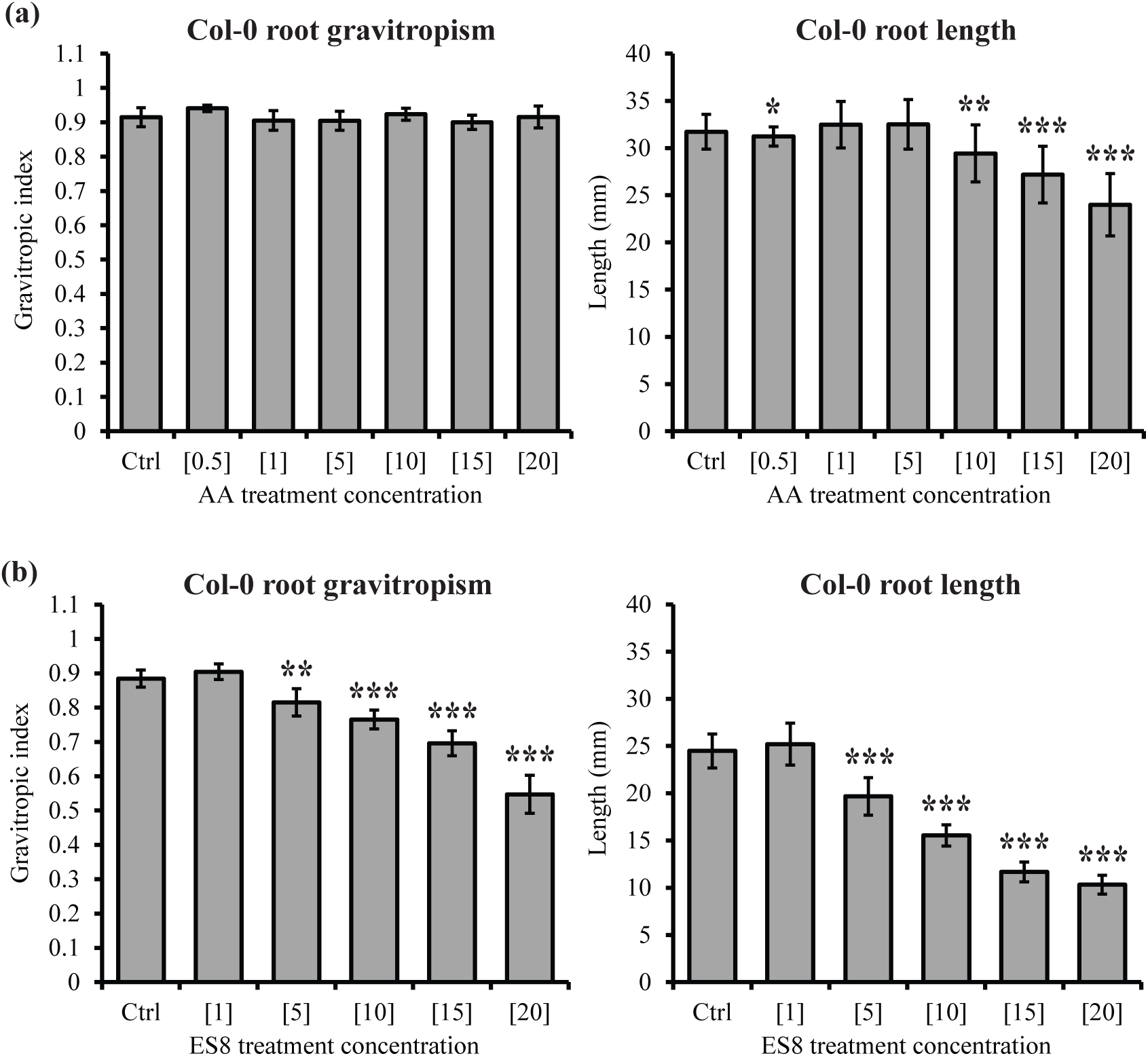
Effects of AA and ES8 on root gravitropic growth and length in the wild type. (a-b) Root gravitropic index and length in 9-d-old seedlings of *Arabidopsis thaliana* Col-0 grown on medium supplemented with a range of concentrations of anthranilic acid (AA) (a) or Endosidin 8 (ES8) (b). Asterisks indicate samples significantly different from the mock-treated control (Ctrl) (***, *P*<0.001; **, *P*<0.01; *, *P*<0.05). Error bars indicate standard error of the mean of the biological replicates. Values in square brackets indicate treatment concentrations in μM. *n* = 25 seedlings per sample per each of three (a) or six (b) biological replicates.

**Fig. S3.**
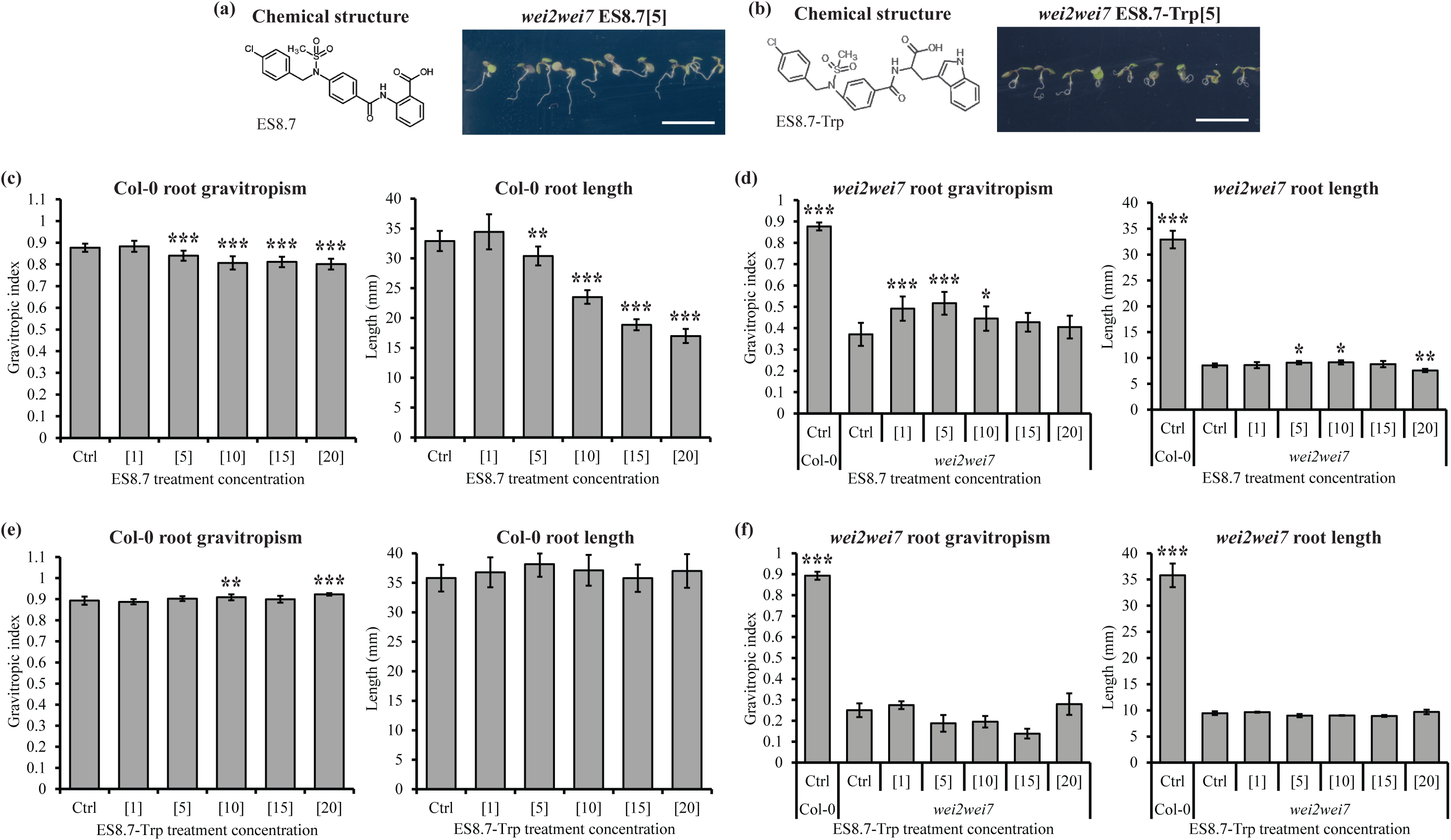
AA but not Trp analogs rescue root gravitropic growth but not length in *wei2wei7*. (a-b) Chemical structures of Endosidin 8.7 (ES8.7) (a) and ES8.7-Tryptophan (Trp) (b) and representative images of 9-d-old *Arabidopsis thaliana wei2wei7* seedlings grown on ES8.7 (a) and ES8.7-Trp (b) -supplemented medium. Scale bars represent 1 cm. (c-f) Root gravitropic index and length in 9-d-old seedlings of *Arabidopsis thaliana* Col-0 (c, e) and *wei2wei7* (d, f) grown on medium supplemented with a range of concentrations of ES8.7 (c, d) or ES8.7-Trp (e, f). Asterisks indicate samples significantly different from the Col-0 mock-treated control (Ctrl) (***, *P*<0.001; **, *P*<0.01; *, *P*<0.05). Error bars indicate standard error of the mean of the biological replicates. Values in square brackets indicate concentrations in μM. *N* = 25 seedlings per samples per each of seven (c, d) or three (e, f) biological replicates.

**Fig. S4.**
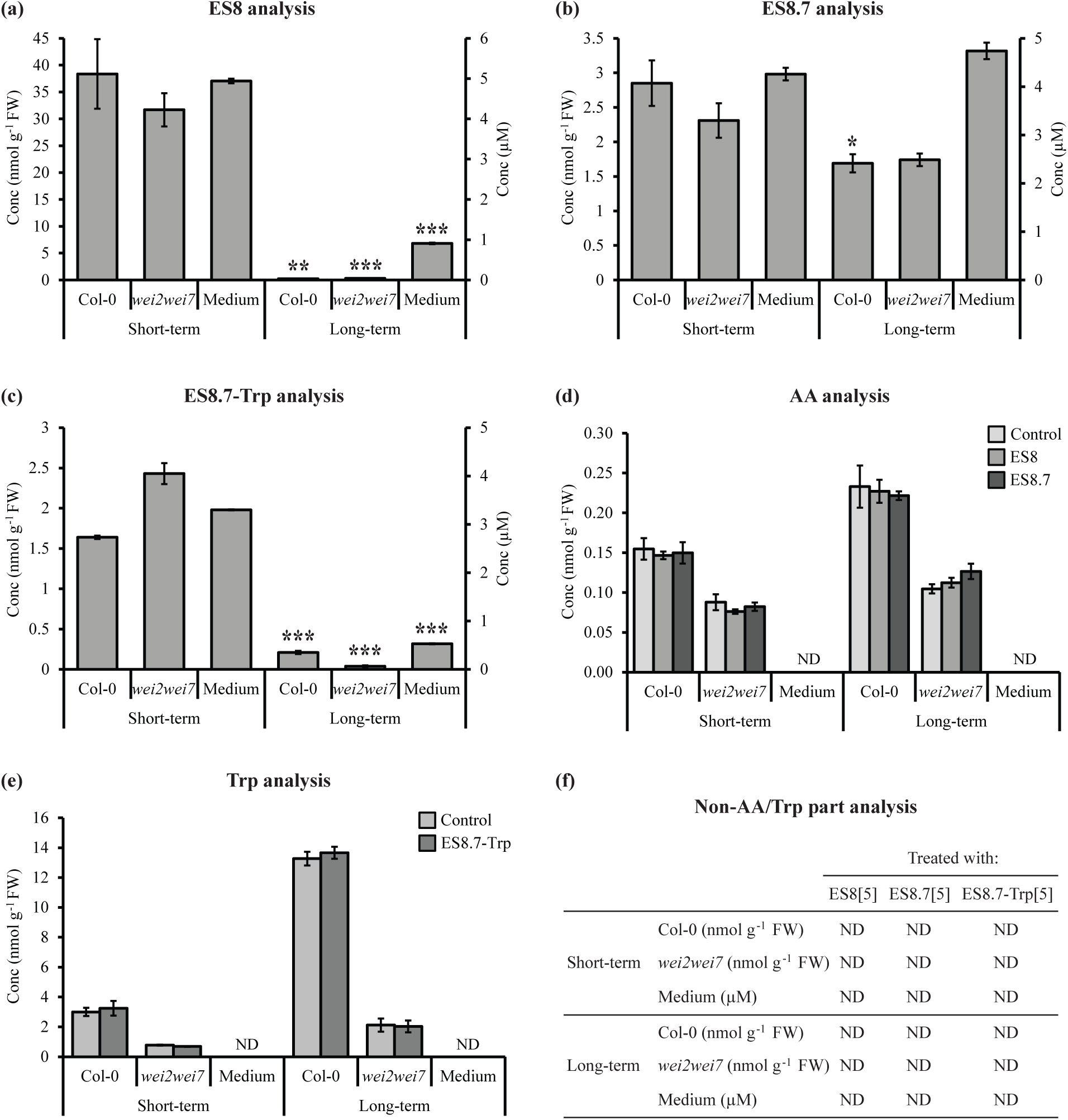
The ES8 compounds are not degraded/metabolized to release AA or Trp. Samples of *Arabidopsis thaliana* Col-0 and *wei2wei7* seedlings after short-term (5-d-old seedlings incubated 5 h in liquid treatment medium) or long-term (9-d-old seedlings grown on solid treatment medium) treatments with ES8 compounds were analyzed for concentrations (Conc) of various compounds. Samples of the short-term and long-term treatment medium, which were incubated for the same length of time but without any seedlings added, were also analyzed. (a) Endosidin 8 (ES8) analysis in ES8-treated samples. (b) ES8.7 analysis in ES8.7-treated samples. (c) ES8.7-Tryptophan (Trp) analysis in ES8.7-Trp-treated samples. (d) Anthranilic acid (AA) analysis after mock treatment (Control) or treatment with ES8 or ES8.7. (e) Trp analysis after mock treatment (Control) or treatment with ES8.7-Trp. (f) Analysis of the non-AA/Trp part of the relevant ES8 compound after treatment with that ES8 compound. Asterisks indicate long-term-treated samples significantly different from the corresponding short-term-treated samples (***, *P*<0.001; **, *P*<0.01; *, *P*<0.05) (a-c). No significant differences were found between ES8/ES8.7/ES8.7-Trp-treated and the corresponding mock-treated control samples (d-e). ND: not detected. Error bars indicate standard error of the mean of the biological replicates. Values in square brackets indicate concentrations in μM. *n* = 100 Col-0 or 250 *wei2wei7* 5-d-old seedlings or 50 Col-0 or 150 *wei2wei7* 9-d-old seedlings per sample per each of two biological replicates, with two technical replicates analyzed per biological replicate.

**Fig. S5.**
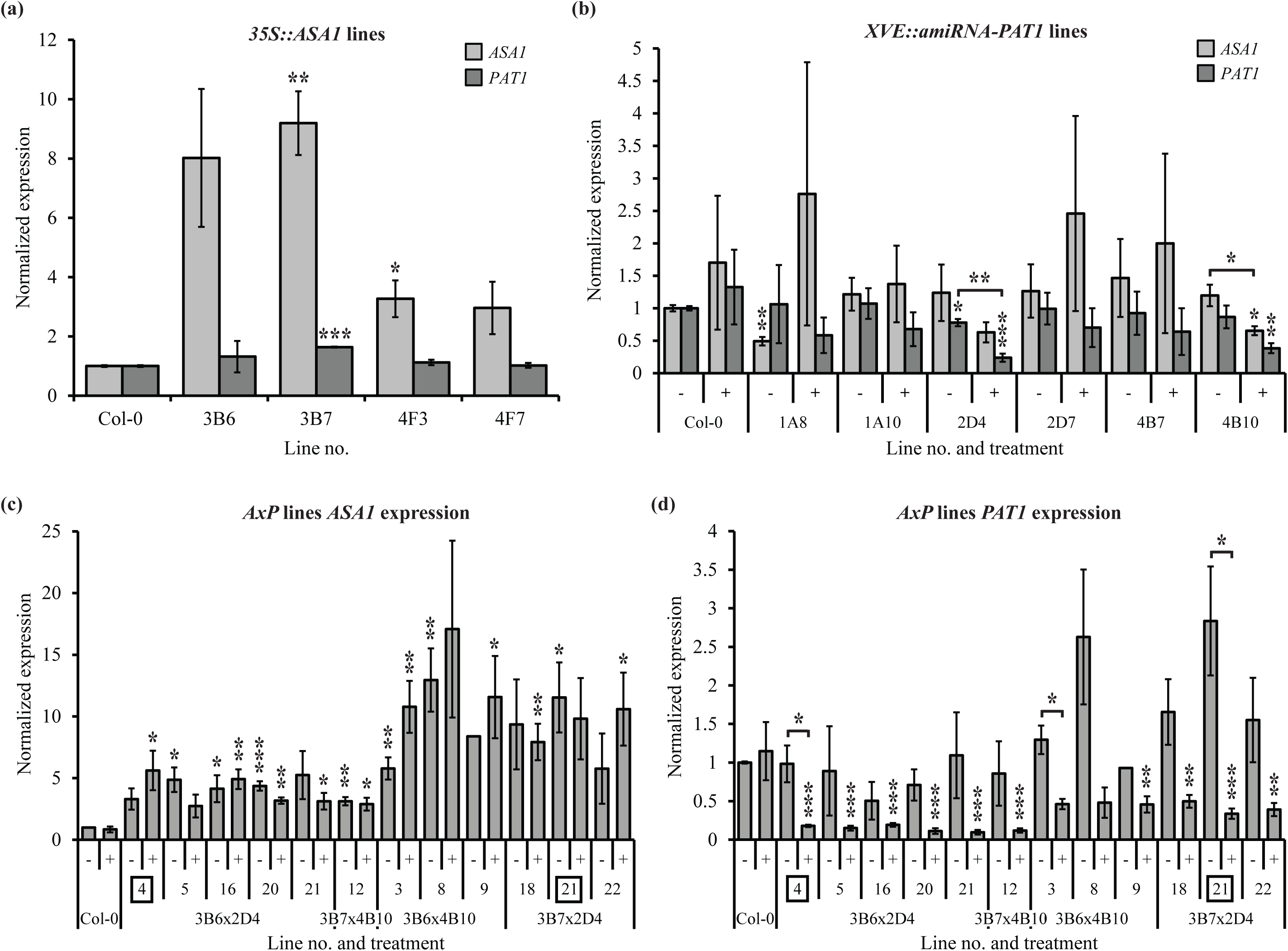
Expression levels of *ASA1* (*WEI2*) and *PAT1* in *AxP* lines. (a) Normalized expression levels of *ASA1* and *PAT1* in 7-d-old *Arabidopsis thaliana 35S::ASA1* seedlings. (b) Normalized expression levels of *ASA1* and *PAT1* in 7-d-old non-induced (-) and estradiol-induced (+) *Arabidopsis thaliana XVE::amiRNA-PAT1* seedlings. (c-d) Normalized expression levels of *ASA1* (c) and *PAT1* (d) in 7-d-old non-induced (-) and estradiol-induced (+) *Arabidopsis thaliana 35S::ASA1 x XVE::amiRNA-PAT1* (*ASA1 x PAT1*; *AxP*) seedlings. The lines marked with squares were selected for further analysis (*AxP1*: 3B6 × 2D4 line no. 4; *AxP2*: 3B7 × 2D4 line no. 21). For estradiol induction of *PAT1* silencing, seedlings were grown on 20 μM estradiol-supplemented medium. Expression values relative to the Col-0 non-induced control are shown. Asterisks indicate samples significantly different from the Col-0 non-induced control unless otherwise indicated (***, *P*<0.001; **, *P*<0.01; *, *P*<0.05). Error bars indicate standard error of the mean of the biological replicates. *n* = 20 seedlings per sample per each of three biological replicates.

**Fig. S6.**
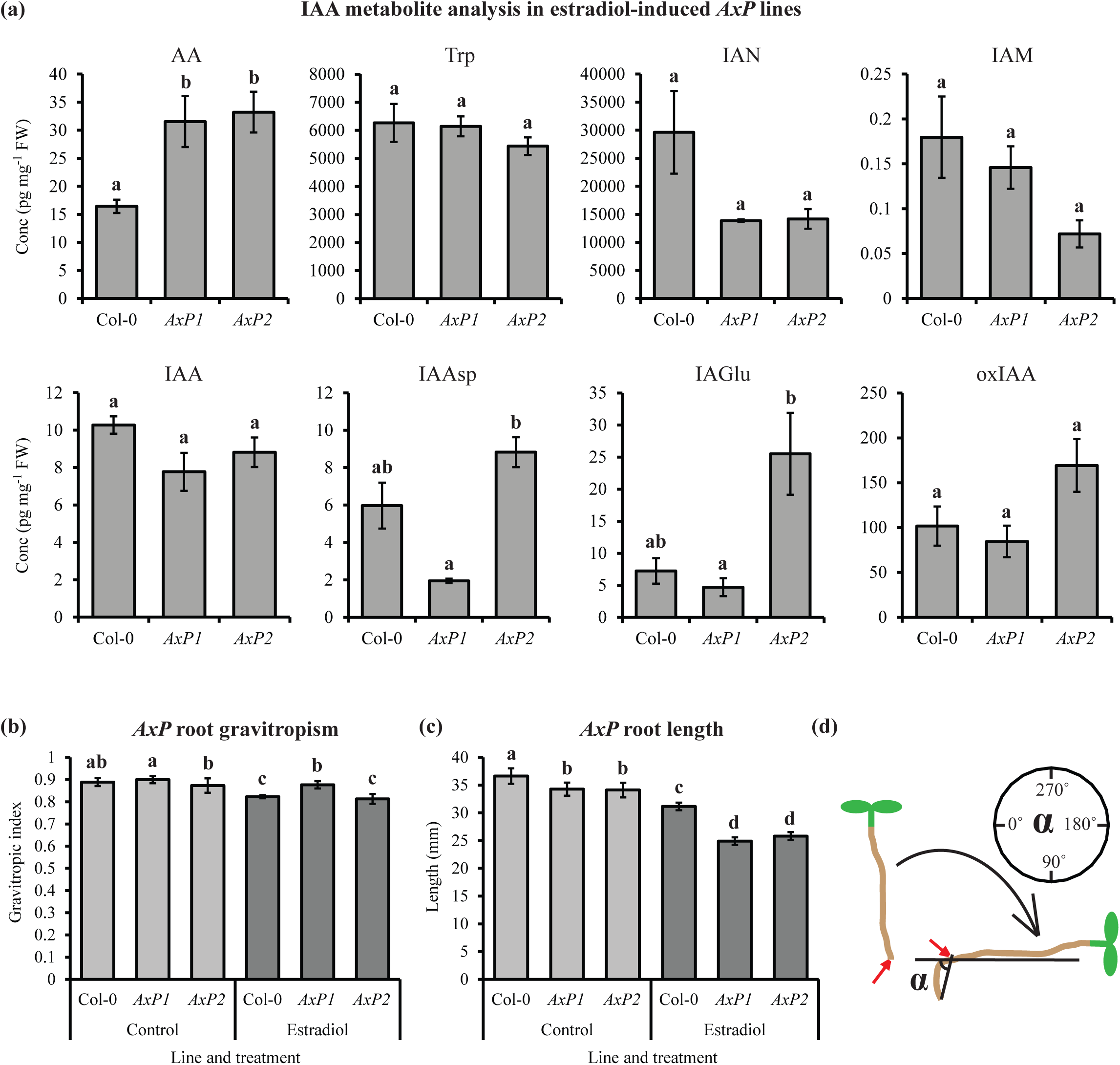
IAA metabolite levels and root phenotypes of *AxP* lines. (a) Concentrations (Conc) of anthranilic acid (AA), tryptophan (Trp), indole-3-acetonitrile (IAN), indole-3-acetamide (IAM), indole-3-acetic acid (IAA), IAA-Aspartate (IAAsp), IAA-Glutamate (IAGlu) and 2-oxoindole-3-acetic acid (oxIAA) in 5-d-old seedlings of *Arabidopsis thaliana* Col-0, *AxP1* and *AxP2* grown on 20 μM estradiol-supplemented medium. (b-c) Root gravitropic index (b) and length (c) in 9-d-old Col-0, *AxP1* and *AxP2* seedlings grown on mock (Control) and 20 μM estradiol-supplemented medium. (d) Scheme representing root bending angle (α) measurements in Col-0 and *AxP* lines after a 90° gravistimulus (black arrow). Angles were measured at the pre-gravistimulus root tip position (red arrows). Different letters indicate significant differences (*P*<0.05). Error bars indicate standard error of the mean of the biological replicates. Tissue was sampled from a mixture of 20 ground seedlings per sample per each of three biological replicates (a). *n* = 20 seedlings per each of three biological replicates (b-c).

**Fig. S7.**
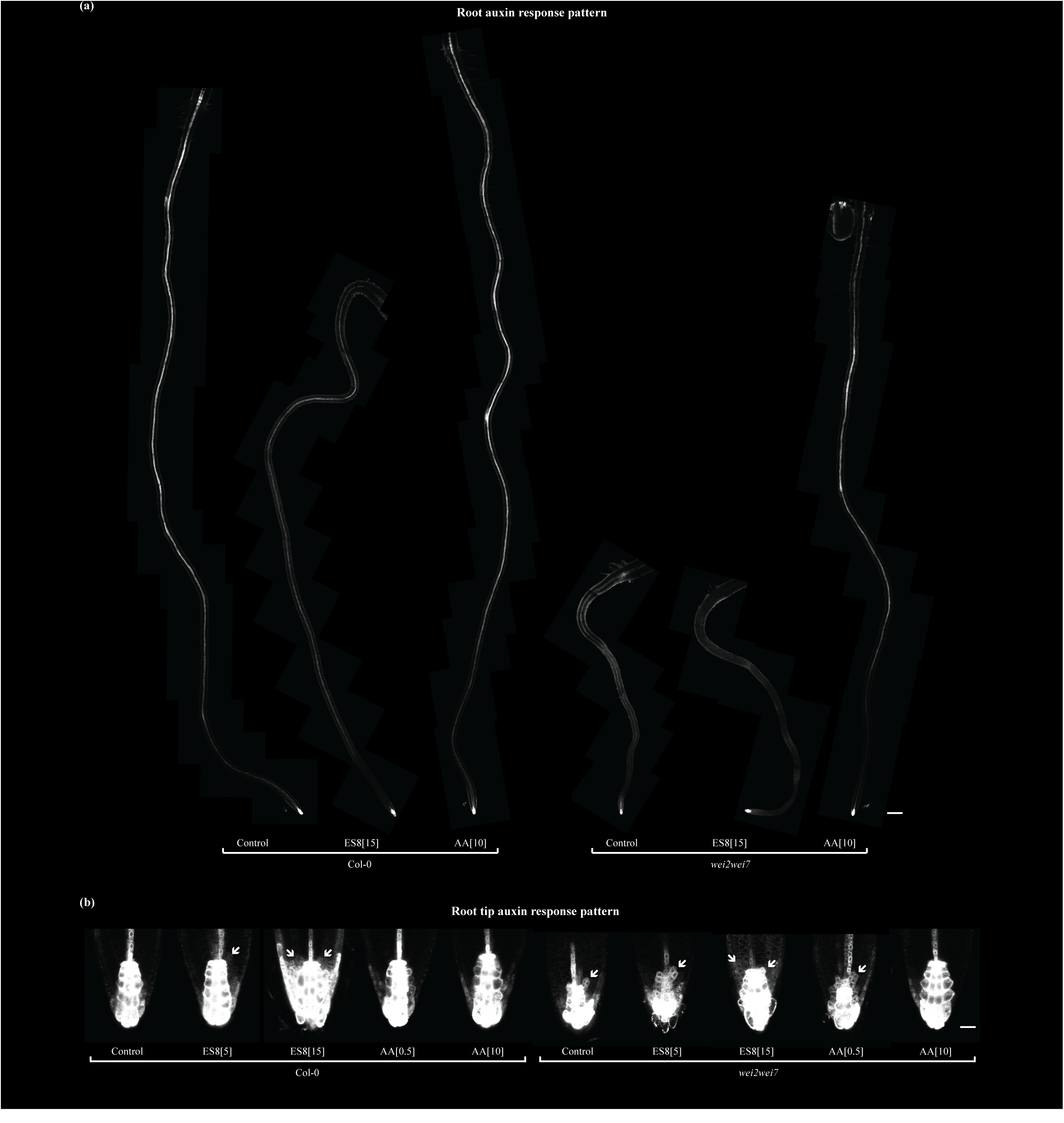
Expression pattern of the auxin-responsive promoter *DR5* is altered in the root by ES8 treatment or AA deficiency. Representative images of GFP fluorescence of *DR5::GFP* in full roots (a) and root tips (b) of 5-d-old *Arabidopsis thaliana* Col-0 and *wei2wei7* seedlings grown on mock (Control), Endosidin 8 (ES8) or anthranilic acid (AA)-supplemented medium. Scale bars represent 200 μm (a) and 20 μm (b). Tiled compilations of images were used to visualize the full roots (a). Accumulations of GFP signal in cell file initials surrounding the quiescent center are marked by white arrows (b). Values in square brackets indicate concentrations in μM. *n* = three (a) or 10 (b) seedlings imaged per sample per each of four (a) or three (b) biological replicates.

**Fig. S8.**
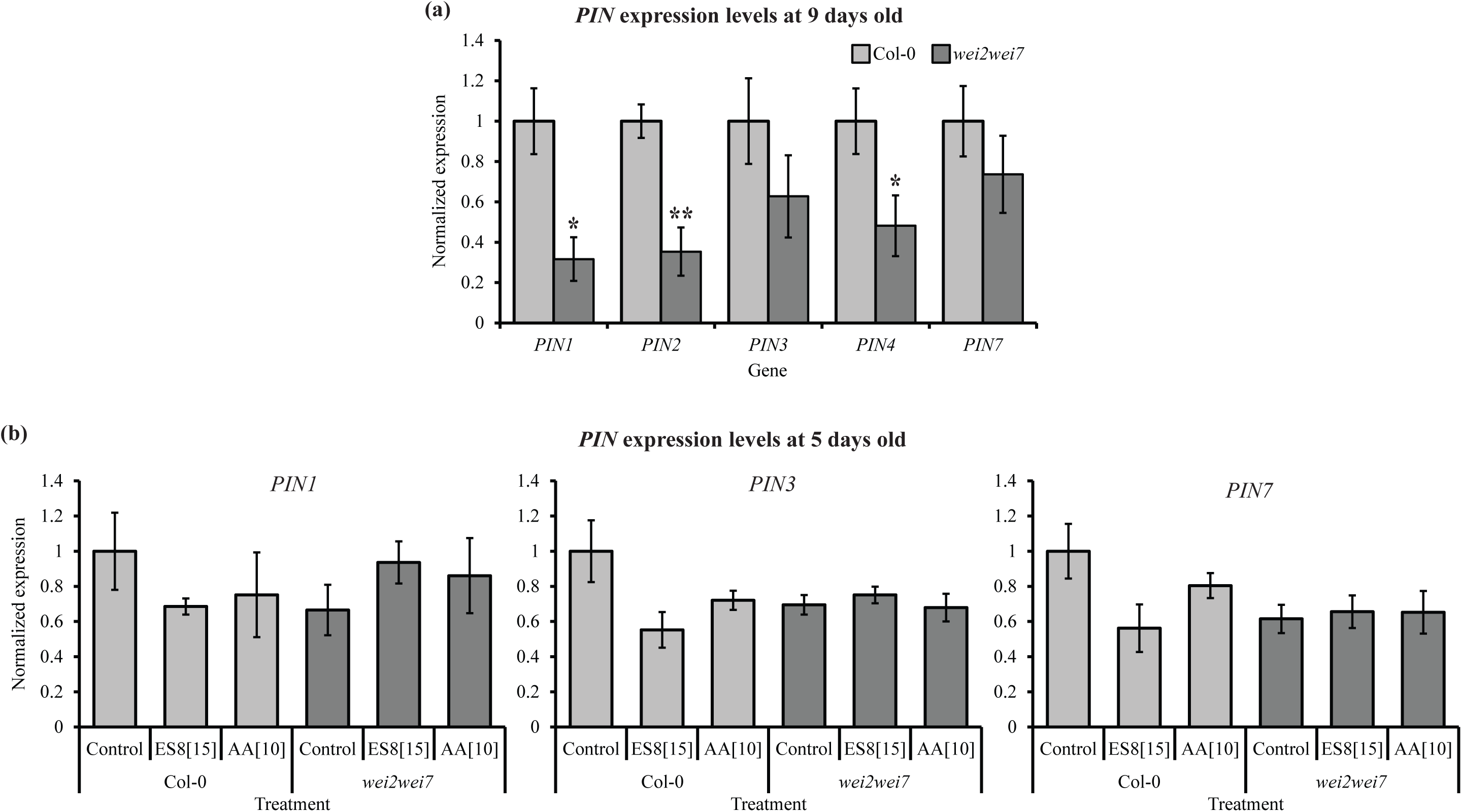
Expression levels of *PIN* genes in Col-0 and *wei2wei7*. (a) Normalized expression levels of *PIN1*, *PIN2*, *PIN3*, *PIN4* and *PIN7* in 9-d-old *Arabidopsis thaliana* Col-0 and *wei2wei7* seedlings. (b) Normalized expression levels of *PIN1*, *PIN3* and *PIN7* in 5-d-old Col-0 and *wei2wei7* seedlings treated with Endosidin 8 (ES8) or anthranilic acid (AA) for 2 h in liquid treatment medium. Expression values relative to the Col-0 control are shown. Graphs show mean of four biological replicates and standard error of the mean. Asterisks indicate significantly different from Col-0 (**, *P*<0.01; *, *P*<0.05). Error bars indicate standard error of the mean of the biological replicates. Values in square brackets indicate concentrations in μM. *n* = 30 seedlings per sample per each of four biological replicates.

**Fig. S9.**
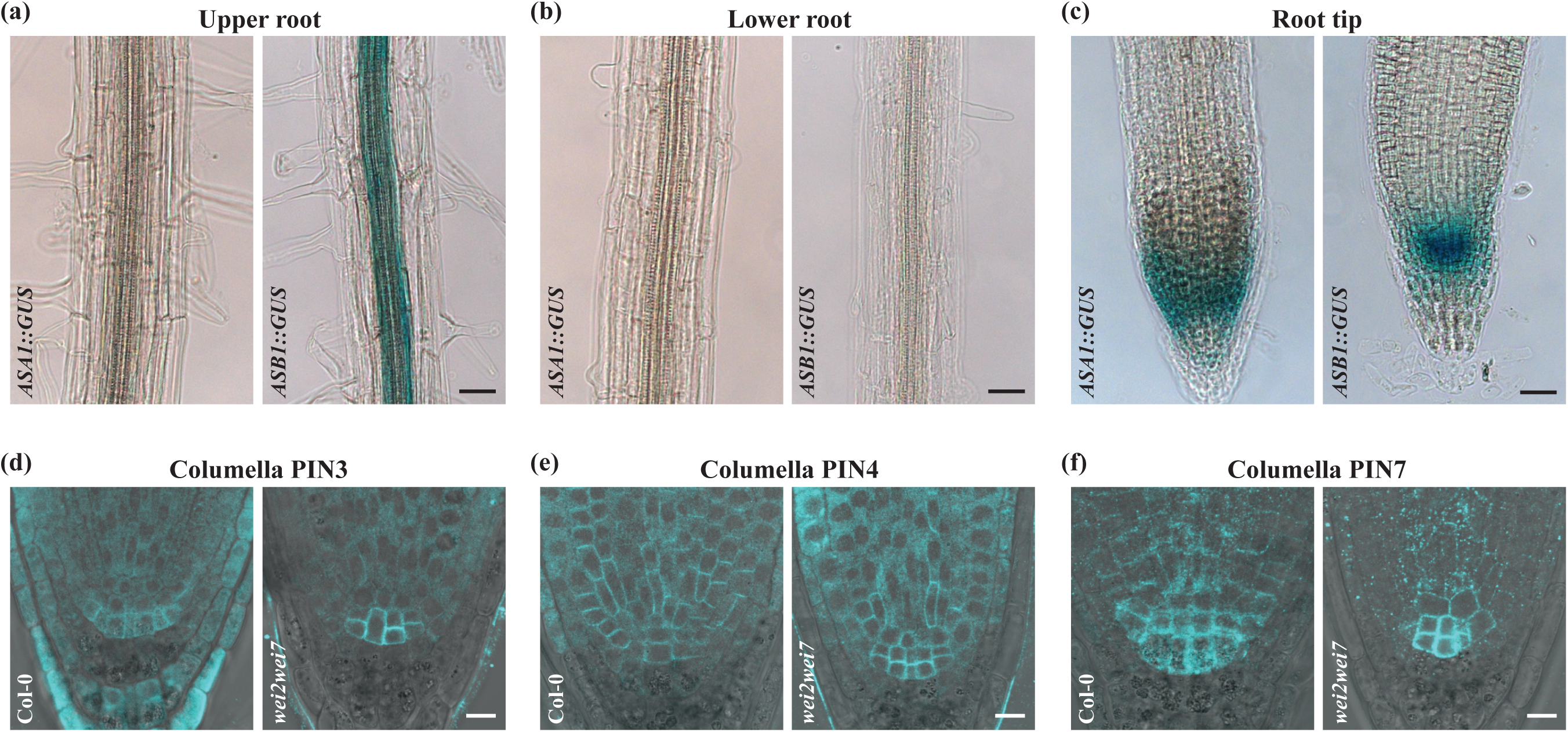
Expression patterns of *ASA1* (*WEI2*) and *ASB1* (*WEI7*) in the root and immunolocalization of PIN3, PIN4 and PIN7 in the columella. (a-c) Representative images of GUS-stained upper roots (a), lower roots (b) and root tips (c) of 9-d-old *Arabidopsis thaliana ASA1::GUS* and *ASB1::GUS* seedlings. Scale bars represent 30 μm. (d-f) Representative images (merged transmission light and Cy3 channel images) of immunolabeled PIN3 (d), PIN4 (e) and PIN7 (f) in root tips of 5-d-old *Arabidopsis thaliana* Col-0 or *wei2wei7* seedlings. Scale bars represent 10 μm. *n* = 10 seedlings imaged per sample per each of three biological replicates.

**Fig. S10.**
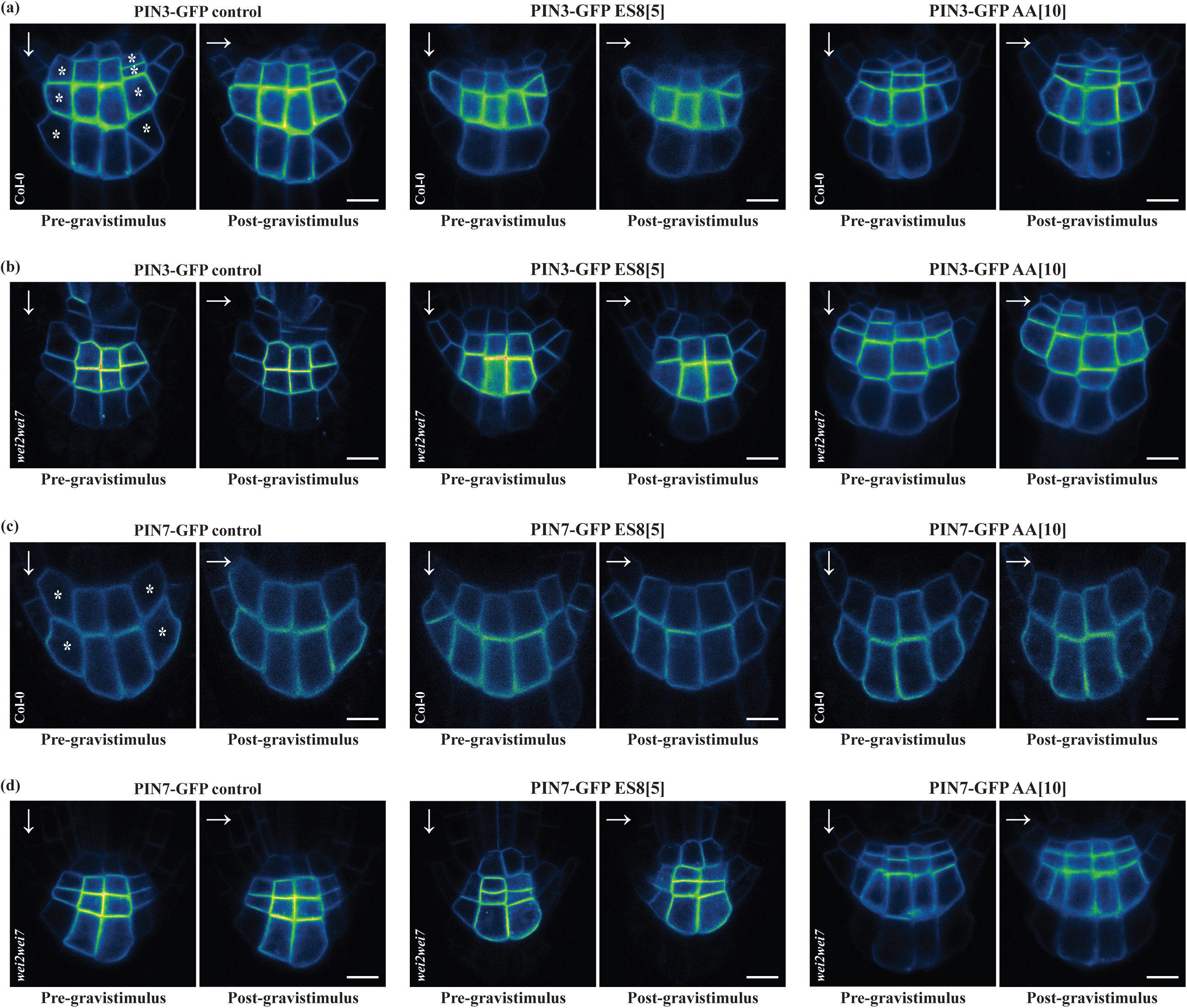
Gravitropic relocalization of PIN3- and PIN7-GFP in root columella cells. Representative images of GFP fluorescence in *PIN3::PIN3-GFP* (a-b) and *PIN7::PIN7-GFP* (c-d) in root columella cells of 5-d-old *Arabidopsis thaliana* Col-0 (a, c) and *wei2wei7* (b, d) seedlings grown on anthranilic acid (AA) and Endosidin 8 (ES8)-supplemented medium directly before and after a 90° gravistimulus of 30 min, during which the seedlings were turned clockwise (the left- and right-facing plasma membranes became upward- and downward-facing, respectively). Arrows represent the direction of the gravity vector immediately prior to image acquisition. Scale bars represent 10 µm. Single confocal plane images are displayed for clarity, but maximal intensity projections of confocal z-stacks were used for the plasma membrane fluorescence analyses displayed in Fig. 5c-f. In the example images where asterisks are shown, these indicate the cells for which fluorescence was measured.

**Fig. S11.**
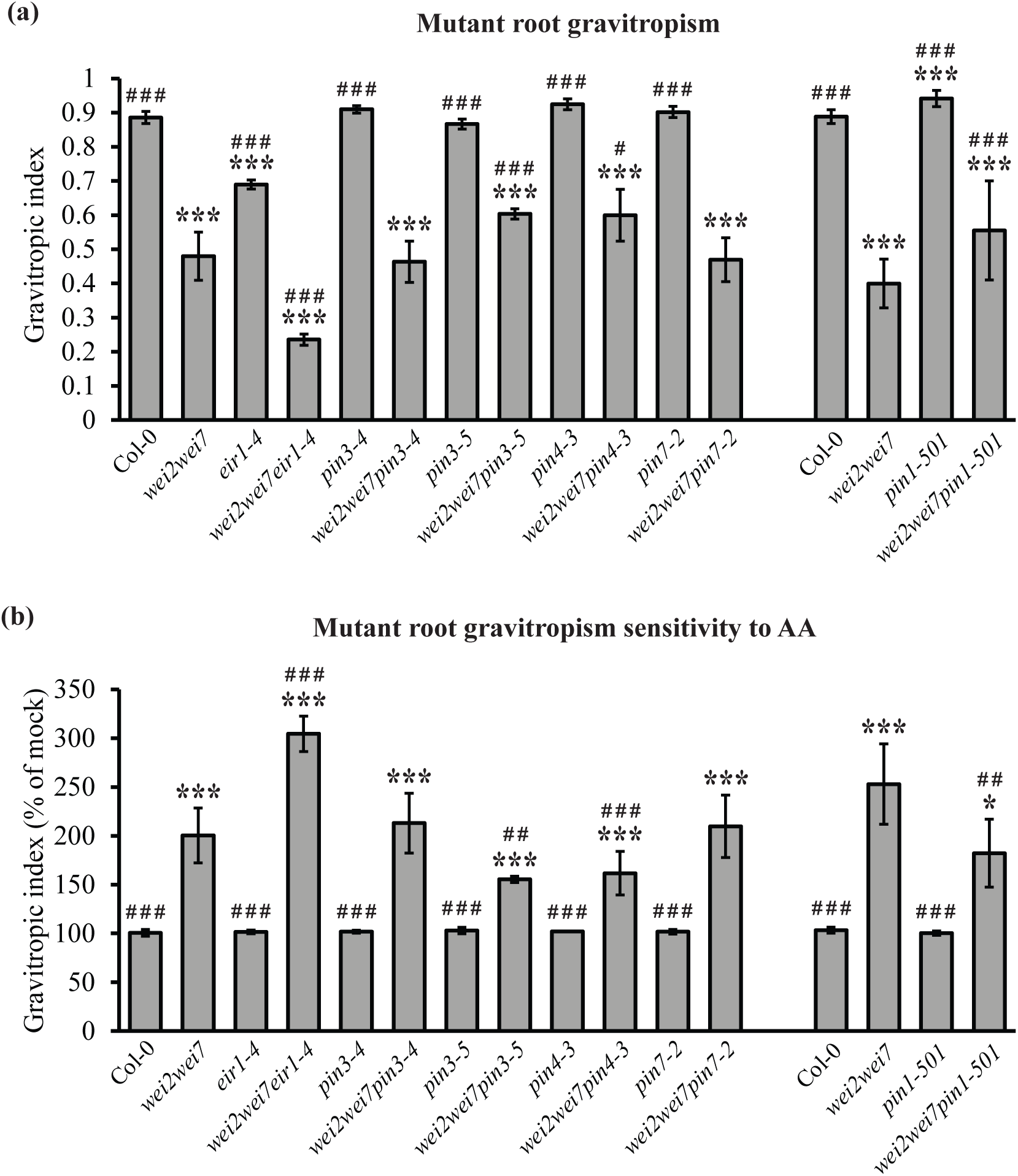
Root gravitropic growth in *pin* and *wei2wei7pin* mutants. (a) Root gravitropic index in 5-d-old seedlings of *Arabidopsis thaliana pin* mutants and *pin* mutants crossed into *wei2wei7*. (b) Root gravitropic index (expressed as a percentage of that in mock-treated samples of the same line) in 5-d-old seedlings of *pin* and *wei2wei7pin* mutants grown on 20 μM AA-supplemented medium. Asterisks and hash symbols indicate significant differences from Col-0 (***, *P*<0.001; *, *P*<0.05) or *wei2wei7* (^# # #^, *P*<0.001; ^# #^, *P*<0.01; ^#^, *P*<0.05), respectively. Error bars indicate standard error of the mean of the biological replicates. *n* = 30 seedlings per sample per each of three biological replicates.

**Fig. S12.**
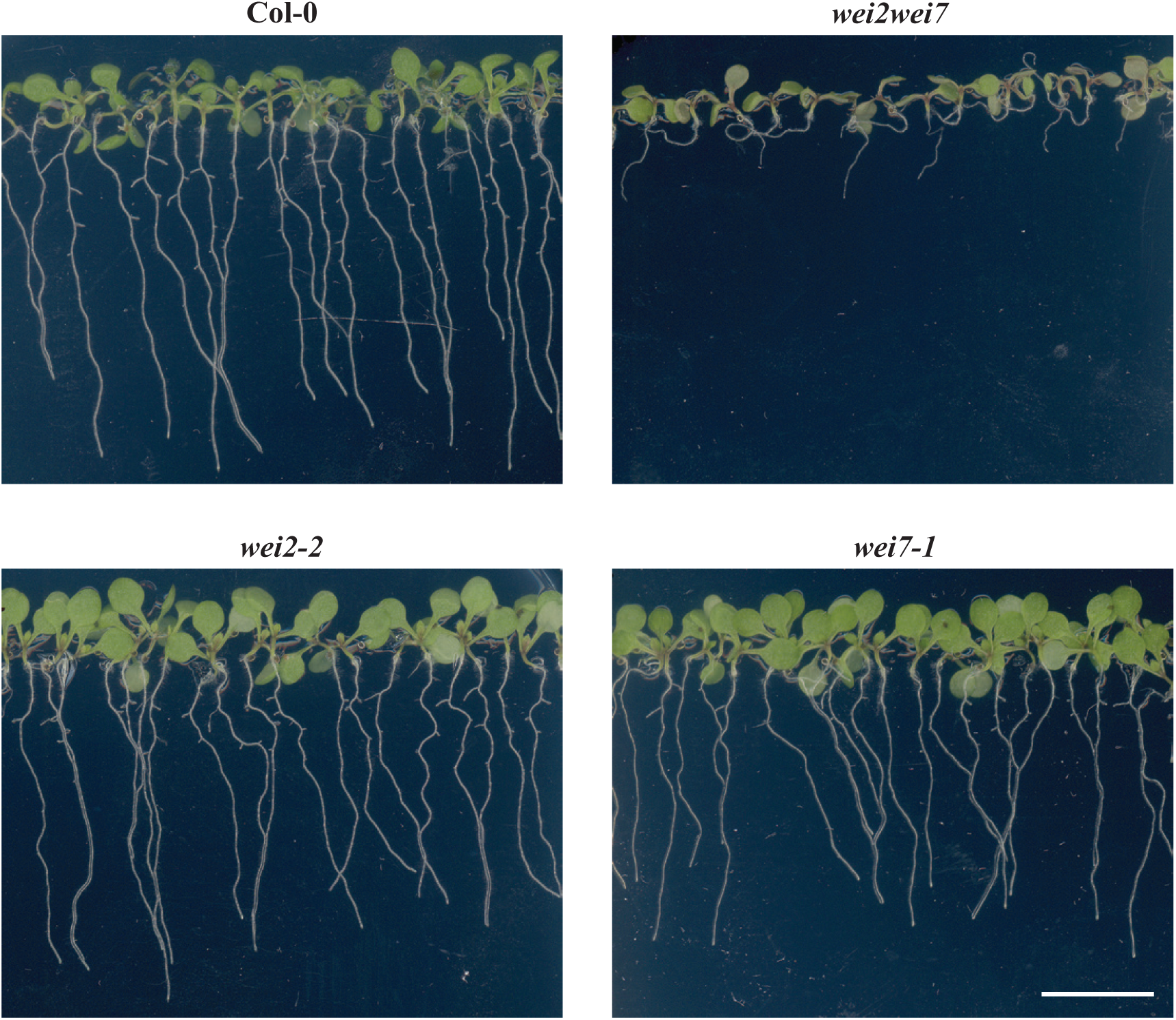
Root phenotypes of double and single mutants of *wei2* and *wei7*. Representative images of 9-d-old A*rabidopsis thaliana* Col-0, *wei2wei7*, *wei2-2* and *wei7-1* seedlings grown on solid medium. Scale bar represents 1 cm.

